# A combinatorial EVs-miRNA signature mediates the anti-tumoral activity of NFAT3-regulated extracellular vesicles in aggressive cancers

**DOI:** 10.64898/2026.02.13.702809

**Authors:** Guénolé Tossou, Nadia Ourari, Maëlle Ralu, Alexandre Montanede, Frédéric Guaddachi, Babette Beher, Manon Paul, Emilie Ouanounou, Morgane le Bras, Stéphane Brunet, Jacqueline Lehmann-Che, Sébastien Jauliac

## Abstract

Aggressive cancers such as triple-negative breast cancer (TNBC) and pancreatic cancer remain difficult to treat because their malignant behavior is driven by complex gene networks rather than single oncogenic targets. Here, we identify an extracellular vesicle (EV)-associated miRNA signature functionally linked to NFAT3 activity and demonstrate its ability to suppress tumor aggressiveness. Functional analyses revealed that a combination of fifteen miRNAs (miR-Comb 15) was required to fully reproduce the anti-tumoral effects of NFAT3-regulated EVs across TNBC and pancreatic cancer models, whereas individual miRNAs showed only partial activity. These effects were associated with coordinated regulation of validated target genes controlling proliferation and invasion, supporting a network-modulating mechanism of action. To facilitate therapeutic translation, we used EVs derived from HEK 293T cells, a non-tumoral, scalable, and readily engineerable EV source. Using an optimized exogenous pH-gradient loading strategy, miR-Comb 15 was efficiently incorporated into EVs without affecting vesicle integrity or intrinsic bioactivity. HEK 293T EVs loaded with miR-Comb 15 consistently showed the strongest anti-tumoral activity in vitro and in vivo among the delivery platforms evaluated. Together, these findings identify a functional NFAT3-dependent EV-miRNA program and support EV-mediated delivery of combinatorial miRNA therapeutics as a promising strategy for aggressive cancers.

## Introduction

Aggressive solid cancers such as Triple-Negative Breast Cancer (TNBC) and pancreatic cancer remain notoriously difficult to treat, largely due to their high proliferative rates, strong invasive behavior, and complex interactions with their microenvironment, making the development of broadly effective treatments particularly challenging^1,2^. Importantly, these aggressive phenotypes arise from the deregulation of multiple interconnected genes and signaling pathways operating within complex oncogenic networks^3^.

In this context, RNA-based therapeutics have emerged as powerful tools for modulating disease-associated gene expression. Among them, miRNAs are uniquely suited to simultaneously regulate multiple interconnected gene networks, making them particularly attractive for counteracting the multifactorial nature of tumor aggressiveness^4^. In this rapidly expanding field of RNA-based therapies, transport remains a major obstacle^5^. Indeed, despite their considerable therapeutic potential, RNA molecules remain vulnerable to degradation, often exhibit suboptimal biodistribution profiles and limited tumor-specific accumulation, and may induce undesirable immune responses^6^. Although synthetic carriers such as lipid nanoparticles have enabled major advances in RNA therapeutics, their clinical use may still be limited by issues related to immunogenicity, tissue tropism, and repeated administration, prompting the development of biological delivery systems such as extracellular vesicles (EVs)^7^. In this context, EVs have emerged as promising natural nanocarriers owing to their intrinsic biocompatibility, ability to transport functional RNA cargo, and role in intercellular communication^8,9^.

EVs are nanoscale lipid bilayer particles released by virtually all cell types and have emerged as potent mediators of intercellular communication. By transporting functional proteins, lipids, and nucleic acids (DNA, mRNA, miRNA, lncRNA and other classes of non-coding RNAs), EVs regulate diverse physiological and pathological processes including tumor progression and immune modulation^10^. Their capacity to protect RNA cargo from degradation and deliver it to recipient cells has generated increasing interest in their exploitation as therapeutic vectors for RNA-based interventions.

Among EVs cargos, microRNAs (miRNAs) represent some of the most functionally impactful. These small non-coding RNAs act as post-transcriptional regulators and are deeply implicated in oncogenesis, metastasis, and therapeutic resistance^11^. Numerous studies have demonstrated their ability to mediate EVs-dependent phenotypes and to orchestrate complex intercellular communication networks^12^. Therefore, harnessing EVs-miRNAs offers a unique opportunity to exploit endogenous regulatory mechanisms for therapeutic benefit. Accordingly, EV-associated miRNAs and engineered EV-based miRNA delivery strategies are increasingly being explored as therapeutic approaches for aggressive solid tumors because of their ability to simultaneously modulate multiple oncogenic pathways involved in tumor progression and metastasis^13^.

NFAT3 (also known as NFATc4) is a member of the Nuclear Factor of Activated T cells (NFAT) family of transcription factors, which regulate diverse physiological processes in a tissue-dependent manner. In particular, NFAT3 has been implicated in neuronal development and survival, where it contributes to the maintenance of neuronal integrity^14^ and resistance to apoptotic stimuli^15^. In addition, NFAT3 participates in the regulation of apoptosis and tissue-specific differentiation programs. These context-dependent physiological functions may partly explain the divergent roles attributed to NFAT3 in different pathological settings, including cancer^16^. While several NFAT family members have been associated by our group and others with pro-tumoral functions^17–25^, our previous studies identified the transcription factor NFAT3 as a strong suppressor of tumor aggressiveness in luminal breast cancer cells^26,27^. We further demonstrated that EVs derived from NFAT3-expressing luminal breast cancer cells can transfer this anti-tumoral activity to highly aggressive cancer models, including TNBC, pancreatic cancer, melanoma, and glioblastoma. Importantly, neither NFAT3 mRNA nor its translated protein was detected within the EVs^28^. This finding suggested that NFAT3 may contribute to the establishment of an anti-tumoral EVs cargo program capable of restraining malignant behaviors across diverse cancer types.

Based on the hypothesis that EVs-associated miRNAs contributed to the anti-tumoral effects of NFAT3-regulated EVs, we performed a comparative miRNA profiling analysis of EVs derived from luminal breast cancer cells. We identified and characterized a specific combination of 15 NFAT3-dependent miRNAs consistently downregulated following NFAT3 depletion. We showed that these 15 miRNAs (named thereafter miR-Comb 15) collectively recapitulate the anti-invasive and anti-proliferative activity previously observed with NFAT3-regulated EVs.

Although T47D-derived EVs enabled the identification of the anti-tumoral miRNA signature described in this study, their malignant origin raises important translational concerns. Tumor-derived EVs may contain oncogenic proteins, lipids, DNA and RNA species, as well as signaling molecules capable of promoting tumor progression, immune modulation, metastatic dissemination and therapy resistance^29–31^. Consequently, despite their utility as a discovery platform, their direct therapeutic application remains questionable. To overcome this bottleneck, we turned to HEK 293T-derived EVs, a scalable non tumoral platform which is widely used and well-established in research and early preclinical studies^32–39^. We loaded the HEK-293T derived EVs with miRNA-Comb 15 utilizing a pH-gradient loading protocol that enables efficient incorporation of miRNAs into EVs while preserving vesicle integrity^40^. Critically, we demonstrated that HEK 293T-derived EVs loaded with miR-Comb 15 consistently displayed the most favorable overall anti-tumoral profile across aggressive cancer models compared with the alternative carriers evaluated. Importantly, functional analyses revealed that miR-Comb 15 coordinately represses gene modules involved in proliferation/cell-cycle progression and invasion/migration/EMT. These results further support the concept that miR-Comb 15 anti-tumoral activity stems from the simultaneous modulation of complementary oncogenic programs rather than a single molecular target.

Collectively, this study establishes a working model and translational framework linking NFAT3 activity to an EV-associated therapeutic miRNA program. By engineering EVs with a rationally designed miRNA combination, our study advances the development of next-generation RNA-based therapies for highly aggressive cancers and establishes a scalable strategy for EVs-mediated miRNA delivery.

## Results

### NFAT3-dependent modulation of EVs-associated miRNAs and functional identification of anti-tumoral miRNA candidates

In earlier work, we established that EVs from T-47D luminal breast cancer cells suppress tumor growth in TNBC models^28^, an effect that fully depends on NFAT3 expression in EVs-secreting cells. Yet, NFAT3 itself is not packaged into the vesicles, neither as mRNA nor protein, indicating that their anti-tumoral activity is mediated by NFAT3-regulated EVs cargo rather than by direct NFAT3 transfer. We therefore sought to uncover these factors mediating the NFAT3-dependent anti-tumor activity of T47D EVs.

Functional delivery of EVs -associated miRNAs remains a matter of debate^41^, although numerous studies have reported that miRNAs are among the molecules most frequently implicated in EVs-associated biological effects and therapeutic potential^42^. We thus hypothesized that miRNAs were key mediators of the NFAT3-dependent anti-tumor activity of T47D EVs and sought to identify them through miRNA profiling.

Accordingly, total miRNAs were extracted from EVs produced by stable T47D cell lines expressing either control shRNA or two independent shRNAs targeting endogenous NFAT3 to achieve stable NFAT3 knockdown. Two independent T47D cell lines expressing distinct shRNAs targeting endogenous NFAT3 were used in this study to minimize the risk of shRNA-specific off-target effects and to increase confidence in the identification of bona fide NFAT3-dependent miRNAs. The efficient and stable knockdown of NFAT3 achieved with these two shRNAs had been previously validated and reported in our earlier study^28^.

Hierarchical clustering of differentially enriched miRNAs in EVs showed that multiple miRNAs were significantly modulated following NFAT3 depletion (Figure 1A and 1B). To increase the robustness of miRNA selection, only miRNAs exhibiting concordant regulation in both shNFAT3 models compared with their respective controls were retained for subsequent analyses. Among the deregulated miRNAs identified in EVs produced by the two NFAT3-silenced cell lines, 21 miRNAs were significantly downregulated and 26 were upregulated. (Figure 1B).

**Figure 1.**
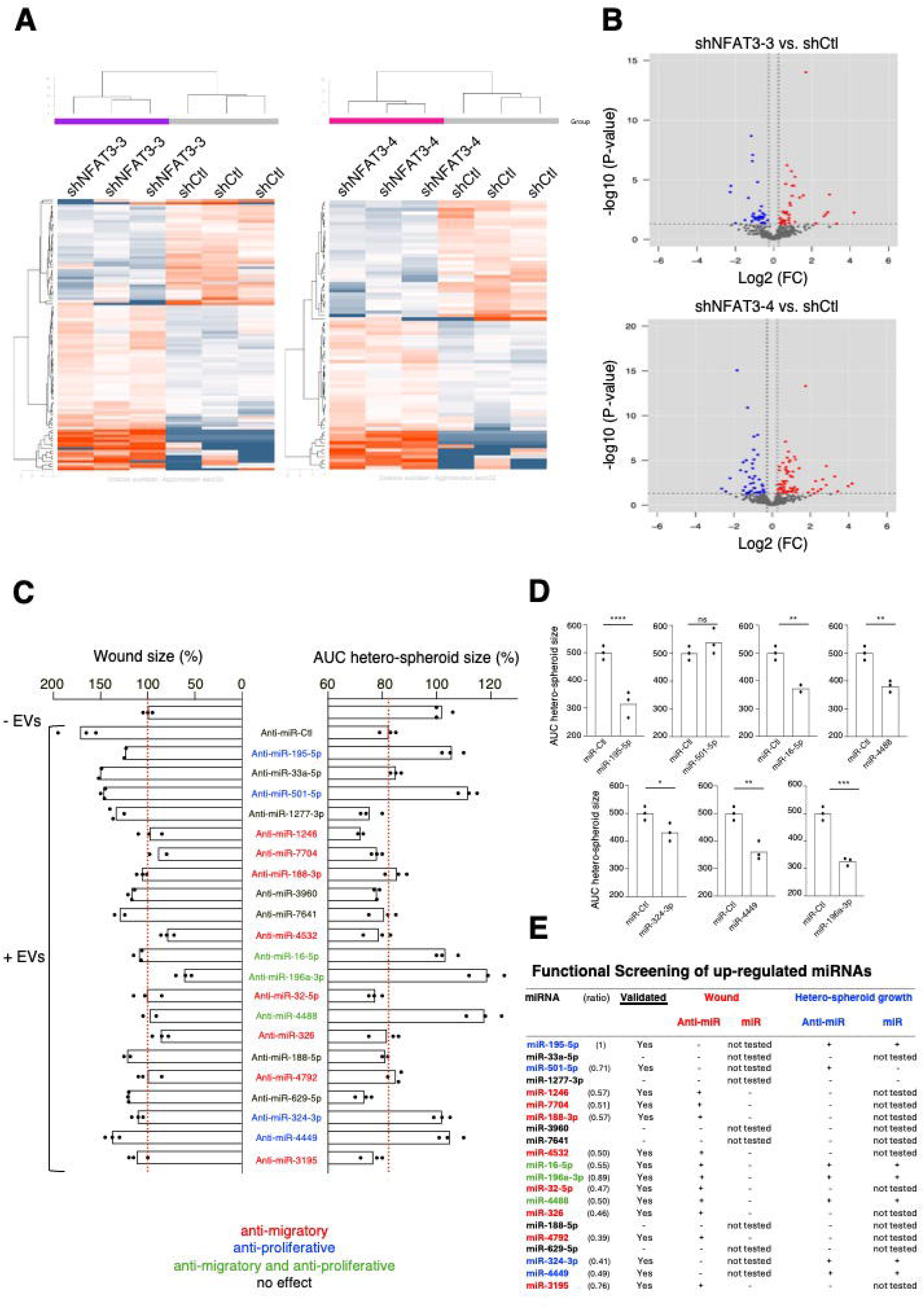
Identification and functional validation of NFAT3-dependent EV-miRNAs mediating anti-tumoral activity. (A) Hierarchical clustering of differentially expressed EV-associated miRNAs from T47D control (shCtl) cells and two independent NFAT3 knockdown clones (shNFAT3-3 and shNFAT3-4). (B) Volcano plots showing differentially expressed miRNAs in EVs from shNFAT3 versus shCtl cells (|log2FC| ≥ 0.25; adjusted *p* ≤ 0.05). (C) Functional screening of individual NFAT3-dependent miRNAs using antagomiRs. Left, wound-healing assay; right, hetero-spheroid growth assay. Red dashed lines indicate the corresponding control values. (D) Validation of selected miRNAs using synthetic miRNA mimics in hetero-spheroid growth assays. (E) Summary of the functional screening, indicating the effects of the antagomir (anti-miRs) and miRNA mimics on migration and hetero-spheroid growth. MiRNA ratios were normalized to miR-195-5p (set to 1) based on miRNA-seq data. Data are presented as mean ± SEM; each dot represents an independent experiment. Statistical significance was determined using an unpaired two-tailed Student’s *t*-test (*ns*, not significant; *P* < 0.05; P < 0.01; *P* < 0.001; P < 0.0001).

We next prioritized the subset of 21 miRNAs that were consistently downregulated following NFAT3 depletion in both independent T47D shNFAT3 cell lines and enriched in EVs derived from NFAT3-expressing cells, as these miRNAs were considered the most likely mediators of the observed anti-tumoral effects.

We functionally evaluated the contribution of individual NFAT3-dependent miRNAs, whose expression was reduced following NFAT3 depletion, to the anti-tumoral activity of EVs by assessing, through a loss-of-function approach, their effects on cell migration using wound-healing assays and on cell proliferation using hetero-spheroid growth assays. We generated hetero-spheroids made of cancer cells and macrophages because we have previously reported that this anti-tumoral action of EVs on cancer cell proliferation requires the presence of macrophages^28^. To this end, we transfected MDA-MB-231 cells with individual antagomirs specific for each miRNA compared to the cells transfected with a control antagomir (sequences in Table S1). These results identified15 of the 21 downregulated miRNAs (miR-Comb 15) as contributors to the anti-tumoral activity of NFAT3-regulated EVs (Figure 1C). Indeed, 8 miRNAs impaired only migration, 4 miRNAs inhibited only proliferation, and 3 miRNAs exhibited both anti-migratory and anti-proliferative activities (Figure 1C and Figures S2, S3). Together, these three classes of miRNAs composing miR-Comb 15 contribute collectively to the anti-tumoral effects mediated by NFAT3-regulated EVs.

To further explore the relationship between NFAT3 activity and the establishment of the miR-Comb 15 signature, we assessed the expression of individual miR-Comb 15 members following NFAT3 depletion in T47D cells. Exploratory RT-qPCR analyses using two independent NFAT3-targeting shRNAs demonstrated that NFAT3 depletion altered the expression of several members of miR-Comb 15 (Figure S1), supporting a functional relationship between NFAT3 activity and the establishment of this anti-tumoral miRNA signature. Combined with the EV RNA-seq data (Figure 1A), these results further suggest that NFAT3 may contribute to the selective enrichment of the miRNA signature into EVs.

Furthermore, we evaluated whether individual transfection of each miRNA into MDA-MB-231 cells was sufficient to inhibit cell migration or cell growth. Six of the seven anti-proliferative miRNAs individually reduced cell growth (Figure 1D). In contrast, none of the identified anti-migratory miRNAs significantly impaired migration when individually transfected (data not shown), suggesting that their coordinated activity is required to suppress migratory behavior. Taken together, our results strongly suggest that miR-Comb 15 is required to fully recapitulate the anti-tumoral effects mediated by NFAT3-regulated EVs (Figure 1E).

### Only the full miR-Comb 15 recapitulates EVs-like anti-tumoral effects across cancer models

To determine whether all 15 miRNAs were required to fully reproduce the anti-tumoral activity of NFAT3-regulated EVs, we generated a reduced miRNA combination (miR-Comb11) containing all miRNAs previously identified as contributing to the anti-migratory phenotype, including the three miRNAs that also displayed anti-proliferative activity (Figure 1A). We then compared the activity of miR-Comb 11 with that of the complete 15-miRNA combination (miR-Comb 15).

The relative proportions of individual miRNAs within miR-Comb 15 were established according to the degree of regulation observed following NFAT3 depletion rather than their endogenous abundance, thereby preserving the hierarchy of NFAT3-dependent miRNA regulation associated with the anti-tumoral EVs program (Figure 1E). Following transfection into two TNBC cell lines (MDA-MB-231 and SUM-159PT) and one pancreatic cancer cell line (MIA PaCa-2), both miR-Comb 11 and miR-Comb 15 significantly inhibited hetero-spheroid growth and invasion compared with miR-Ctl. However, miR-Comb 15 consistently produced a significantly stronger anti-tumoral effect than miR-Comb11 in all three models (Figure 2A), demonstrating that the complete 15-miRNA combination is required to fully recapitulate the anti-proliferative and anti-invasive activities previously observed with NFAT3-regulated EVs. Alterations in cell size have been shown to negatively impact tumor cell proliferation and invasive capacity^43,44^. Thus, we investigated whether direct transfection of miR-Comb 15 would induce detectable cell morphological changes by immunofluorescence staining. miR-Comb 15 transfection was performed in MDA-MB-231, SUM-159PT and MIA-PaCa-2 cell lines. The clustered growth pattern exhibited by SUM-159PT cells prevented accurate single-cell morphometric analyses under our experimental conditions. In the two other cell lines, miR-Comb 15 transfection resulted in a significant increase in cell area compared with miR-Ctl-transfected cells (Figure 2B). Importantly, because miR-Comb 15-treated cells occupy a larger surface area, the concomitant reduction in hetero-spheroid size observed in our functional assays is unlikely to reflect simple morphological alterations. Instead, smaller spheroids despite larger individual cell size most likely indicate a decrease in the overall number of cells constituting the spheroids. This technical limitation was restricted to cell area measurements and did not affect the functional assays performed in SUM-159PT cells, which consistently demonstrated anti-tumoral effects of miR-Comb 15 in this model.

**Figure 2.**
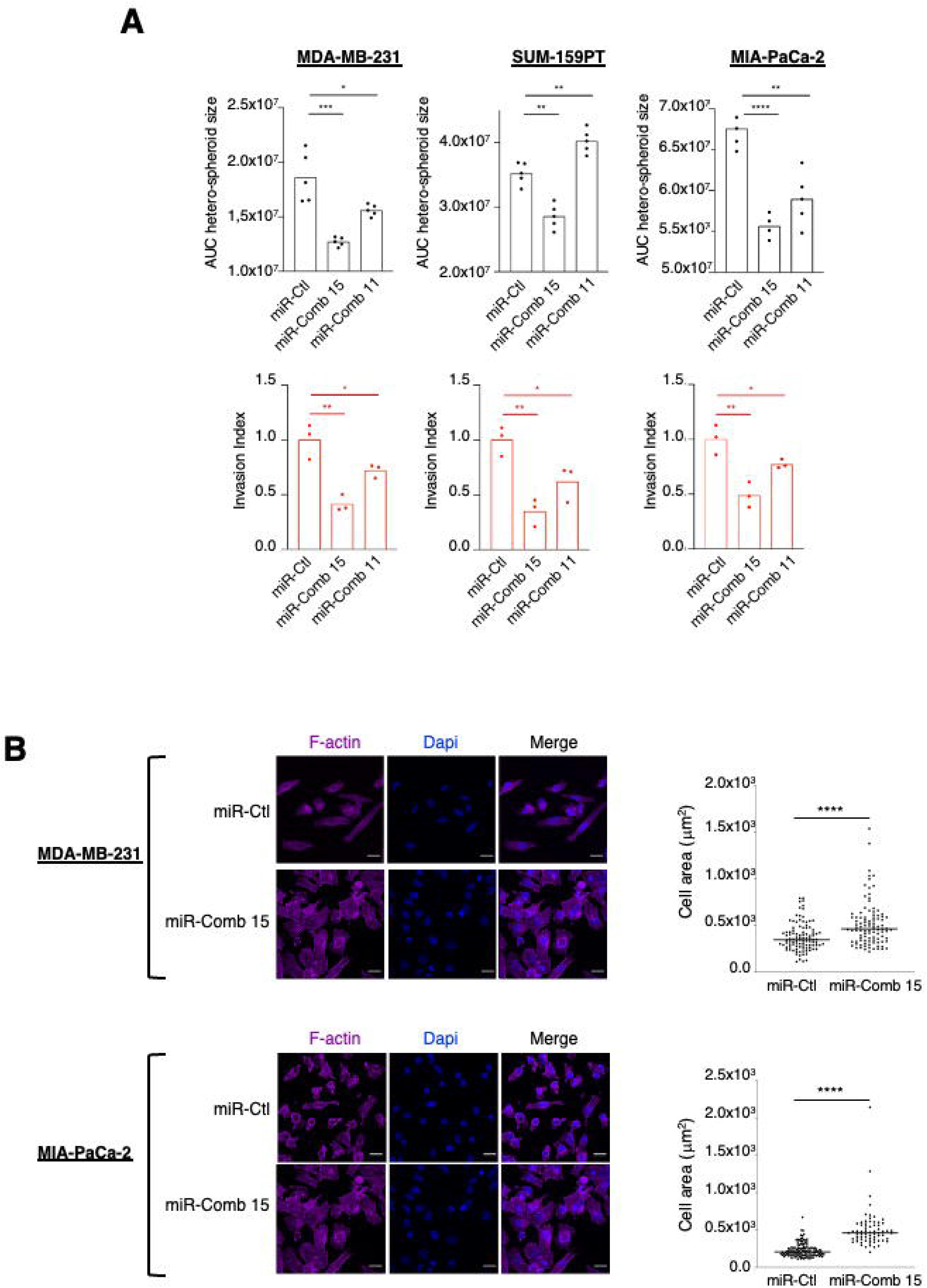
Functional validation of miRNA combinations reproducing the anti-tumoral activity of NFAT3-regulated EVs. (A) Effects of miR-Comb 15 and the reduced miR-Comb 11 on hetero-spheroid growth (upper panels) and invasion (lower panels) in MDA-MB-231, SUM-159PT, and MIA PaCa-2 cells. (B) Representative immunofluorescence images of MDA-MB-231 and MIA PaCa-2 cells transfected with miR-Ctl or miR-Comb 15 and stained for F-actin (phalloidin, magenta) and nuclei (DAPI, blue). Right, quantification of cell area. Scale bars, 20 μm. Data are presented as mean. Each dot in (A) represents an independent experiment (n = 6 for hetero-spheroids assay and n = 3 for invasion assay); each dot in (B) represents a single cell. Statistical significance was determined using an unpaired two-tailed Student’s *t*-test (*ns*, not significant; *P* < 0.05; P < 0.01; *P* < 0.001; P < 0.0001 versus miR-Ctl).

Altogether, these results show that only miR-Comb 15 fully recapitulates the anti-tumoral effects of the EVs, including those acting solely on proliferation being reproducibly required across three aggressive breast and pancreatic cancer cell lines. Based on these findings, all subsequent experiments in this study were conducted using miR-Comb 15.

### miR-Comb 15 modulates oncogenic networks associated with tumor progression

To gain insight into the molecular mechanisms underlying the anti-tumoral activity of miR-Comb 15, experimentally validated targets were retrieved from TarBase and subjected to functional enrichment analyses (https://dianalab.e-ce.uth.gr/tarbasev9)^45^. The resulting list of experimentally validated targets was subjected to Gene Ontology enrichment analyses. These analyses revealed a significant enrichment of biological processes related to cell-cycle progression and proliferative signaling, including positive regulation of cell cycle, epithelial cell proliferation, stem cell proliferation, and positive regulation of growth, as well as processes associated with tumor dissemination, such as regulation of cytoskeleton organization, cell adhesion, cell–substrate adhesion, and cell motility (Figure 3A).

**Figure 3.**
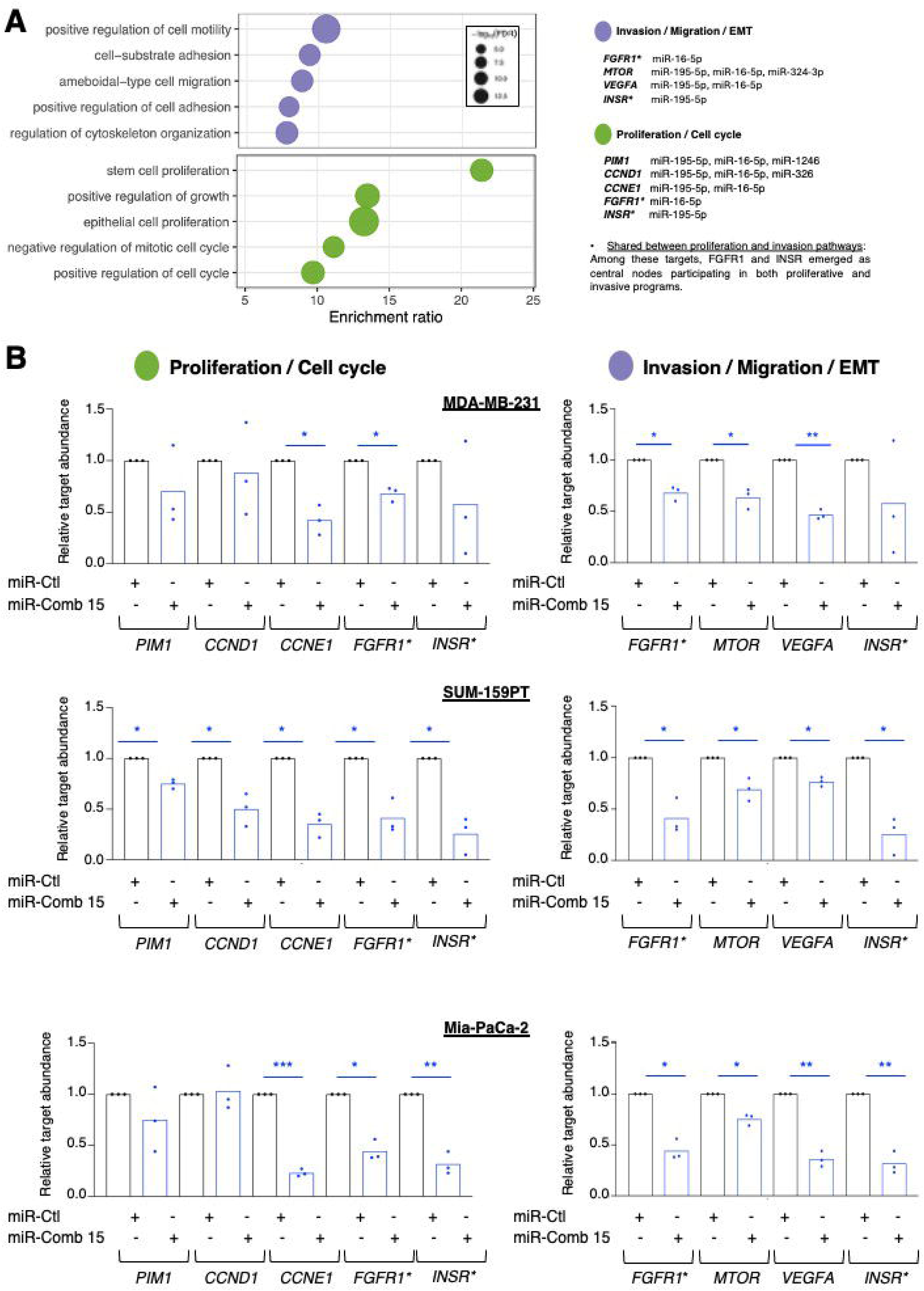
Identification and validation of miR-Comb 15-regulated targets involved in proliferation and invasion programs. (A) Functional enrichment analysis of experimentally validated miR-Comb 15 targets identified using TarBase. Enriched biological processes associated with cell motility, migration, adhesion, cytoskeleton organization, proliferation, and cell cycle regulation are shown. Selected TarBase-validated miRNA targets illustrating the invasion/migration/EMT and proliferation/cell-cycle functional modules are shown.. *FGFR1* and *INSR* emerged as shared targets linking proliferative and invasive programs. (B) RT-qPCR validation of selected miR-Comb 15 target genes in MDA-MB-231, SUM-159PT, and MIA PaCa-2 cells transfected with miR-Comb 15 or miR-Ctl. Relative target abundance was determined by RT-qPCR and normalized to miR-Ctl-transfected cells (set to 1). Genes associated with proliferation/cell cycle pathways (*PIM1*, *CCND1*, *CCNE1*, *FGFR1,* and INSR) and invasion/migration/EMT pathways (*FGFR1*, *MTOR*, *VEGFA*, and *INSR*) are shown. Bars represent the mean of three independent biological experiments (n = 3), with individual data points shown. Statistical significance was determined using Student’s *t*-test. *P* < 0.05; \**P* < 0.01; \*\**P* < 0.001.

To experimentally validate these bioinformatic predictions, we quantified the expression of representative target genes involved in these oncogenic programs following transient transfection with miR-Comb 15 in the three model cell lines used in this study. RT-qPCR analyses demonstrated that miR-Comb 15 reduced the expression of several genes implicated in cell-cycle progression and proliferation, including *PIM1*, *CCND1*, *CCNE1*, *FGFR1*, and *INSR*, across TNBC (MDA-MB-231 and SUM-159PT) and pancreatic cancer (MIA PaCa-2) cell models, although the magnitude of these effects varied depending on the cellular context (Figure 3B). Likewise, miR-Comb 15 decreased the expression of genes associated with invasion and metastatic dissemination, including *FGFR1*, *MTOR*, *VEGFA*, and *INSR*, several of which were significantly modulated in all three cellular models examined. Notably, FGFR1 and INSR were identified as shared nodes between proliferative and invasive pathways, supporting the notion that miR-Comb 15 can simultaneously modulated multiple hallmarks of tumor aggressiveness.

Collectively, these findings support the notion that the anti-tumoral effects induced by miR-Comb 15 arise from the coordinated modulation of interconnected oncogenic networks rather than from the inhibition of a single molecular target. These results provide additional support for the concept of network modulation by miR-Comb 15 and may contribute to explaining its broad anti-tumoral activity across distinct aggressive cancer models.

### Addressing oncogenic concerns of T47D EVs through replacement with HEK 293T EVs

Since T47D cells are of cancerous origin, there is a possibility that their EVs cargo may contain undesirable or oncogenic components. If a potential therapeutic application is envisioned, a more suitable and safer EVs source will be required. To overcome this bottleneck, we selected HEK 293T cells as an alternative EVs-producing platform. Knowing that these cells do express NFAT3^28^, we previously demonstrated that their EVs themselves exert potent anti-tumoral effects in vitro, thereby ensuring that the therapeutic activity observed with T47D-derived EVs would be preserved^28^.

To rigorously identify the optimal dose of these EVs to impede cell growth and invasion, we first performed in vitro dose-response experiments in the three aggressive cancer cell lines used in this study for hetero-spheroids growth assays. Although we tested EVs concentrations higher than 3×10⁸ pp/mL, no improvement in inhibition of hetero-spheroids growth was observed (data not shown). Surprisingly, a lower dose of 3×10³ particles emerged as the optimal concentration for growth inhibition (Figure 4A, left panel). Consistently, no anti-proliferative effect was detected in the SUM-159-PT hetero-spheroids growth assay. We next validated this concentration in invasion assays and confirmed that 3×10³ pp/mL represented the optimal dose for both hetero-spheroid growth and invasion assays (Figure 4A). These findings define the effective dosing parameters for HEK 293T-derived EVs and establish the conditions applied in the subsequent in vitro experiments.

**Figure 4.**
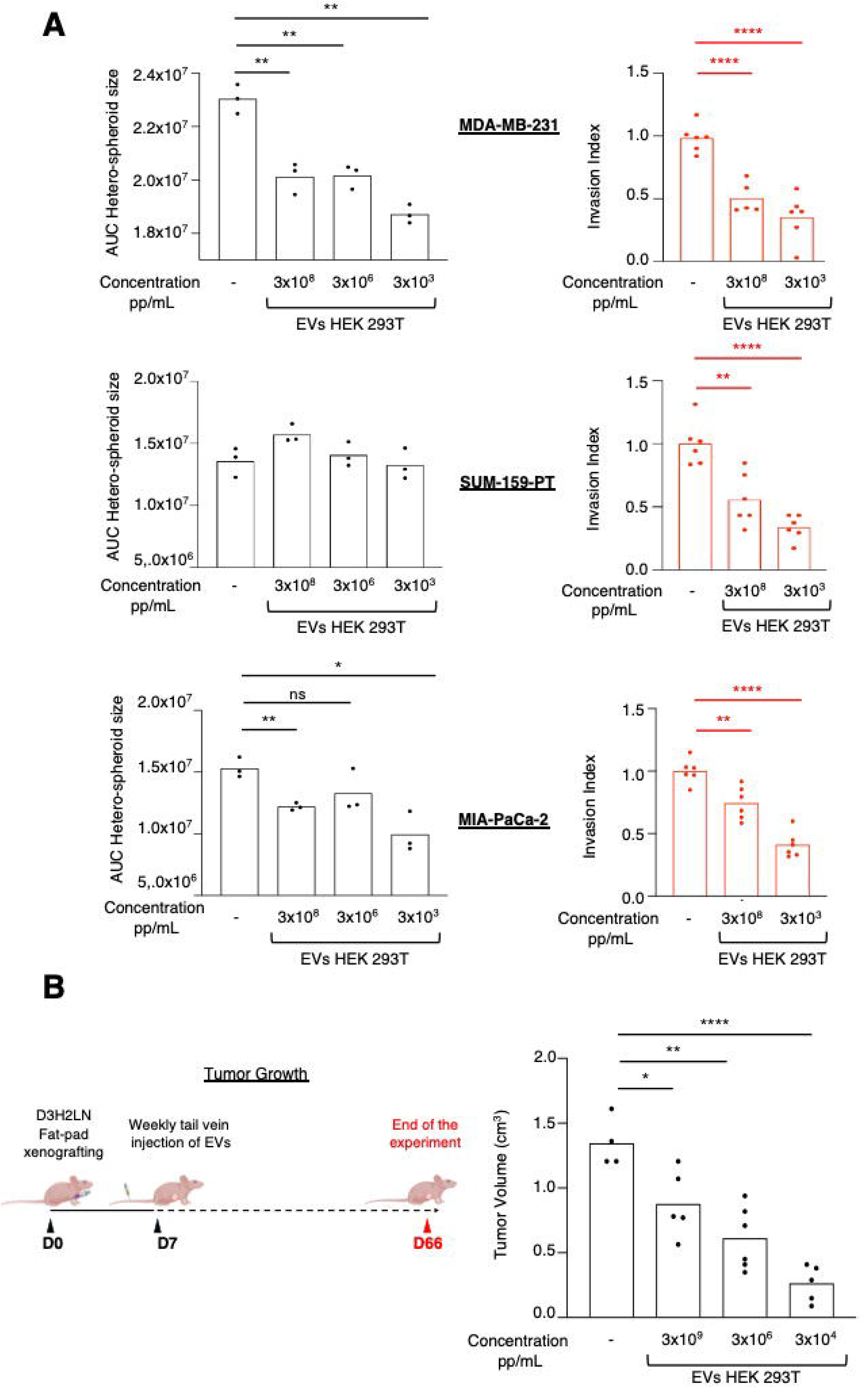
Replacement of T47D-derived EVs by HEK 293T-derived EVs and dose optimization in vitro and in vivo. (A) Dose-response analysis of HEK 293T-derived EVs on hetero-spheroid growth (left, n=3) and invasion (right, n = 6) in MDA-MB-231, SUM-159PT, and MIA PaCa-2 cells. Hetero-spheroid growth was quantified as the area under the curve (AUC). Invasion assays were performed using low (3× 10³ pp/mL) and high (3×10⁸ pp/mL) EVs concentrations. (B) In vivo dose optimization using an orthotopic D3H2LN TNBC mouse model. Left, experimental design. Right, tumor volume measured at day 66 after weekly intravenous administration of PBS or HEK 293T-derived EVs at the indicated doses. Data are presented as mean. Each dot represents an independent experiment (A) or one mouse (B, n = 6 mice per condition). Statistical significance was determined using an unpaired two-tailed Student’s *t*-test (*ns*, not significant; *P* < 0.05; P < 0.01; P < 0.0001 versus untreated or PBS-treated controls).

We then evaluated whether this dosing strategy validated in vitro could be reliably extrapolated to our in vivo model to determine the effective therapeutic dose. As only HEK 293T-ΔNFAT3 EVs^28^, but not unmodified HEK 293T EVs, suppressed tumor growth in vivo (Figure S4), all in vivo dosing experiments were performed exclusively with HEK 293T-ΔNFAT3–derived EVs. Using our established orthotopic TNBC model^28^, athymic nude mice received weekly intravenous injections of PBS or several concentrations of HEK 293T-ΔNFAT3 EVs beginning one week after tumor cells implantation. The highest concentration corresponded to that used in our previous study^28^ and served as the upper reference (Figure 3B). Tumor growth was monitored by caliper measurements. Although all EVs doses significantly inhibited tumor growth, the 3×10⁴ pp/mL yielded the strongest suppression.

These results define the EVs concentration of 3×10⁴ pp/mL as the working dose for subsequent in vivo investigations.

### Efficient exogenous loading of miRNA Comb-15 potentiates the intrinsic anti-tumoral activity of HEK 293T-derived EVs

Having demonstrated that direct transfection of miR-Comb 15 exerts potent anti-tumoral effects in aggressive cancer cell models, we next investigated whether HEK 293T-derived extracellular vesicles (EVs) loaded with this miRNA combination could reproduce and enhance these inhibitory effects. Because HEK 293T-derived EVs were used as carriers for miR-Comb 15, we first characterized the endogenous abundance of each member of this miRNA signature in naïve HEK 293T-derived EVs. RT-qPCR analyses revealed heterogeneous expression levels among the different miRNAs (Figure S5). While some miRNAs were readily detectable, others were present at very low levels or close to the assay detection limit, indicating that endogenous HEK 293T-derived EVs do not naturally recapitulate the complete miR-Comb 15 signature.

To generate EVs enriched in miR-Comb 15, we employed a previously described pH gradient-mediated loading strategy^40^, which enables efficient incorporation of small RNAs while minimizing vesicle disruption and aggregation.

To determine the optimal loading conditions, HEK 293T-derived EVs were incubated with increasing concentrations (200–500 nM) of miR-Comb 15 or a miRIDIAN Negative Control (miR-Ctl). EV-associated miRNAs were subsequently extracted and quantified by RT-qPCR using specific primers for each member of the miRNA combination. As shown in Figure 5A and Figure S6, increasing miRNA concentrations resulted in progressively higher levels of EVs-associated miRNAs up to approximately 400 nM. Beyond this concentration, a plateau or slight decrease was observed, suggesting saturation of the loading capacity under the experimental conditions employed. Importantly, EVs size distributions remained comparable across all loading conditions (Figure S7), indicating that the pH-gradient procedure did not alter EVs physical properties.

**Figure 5.**
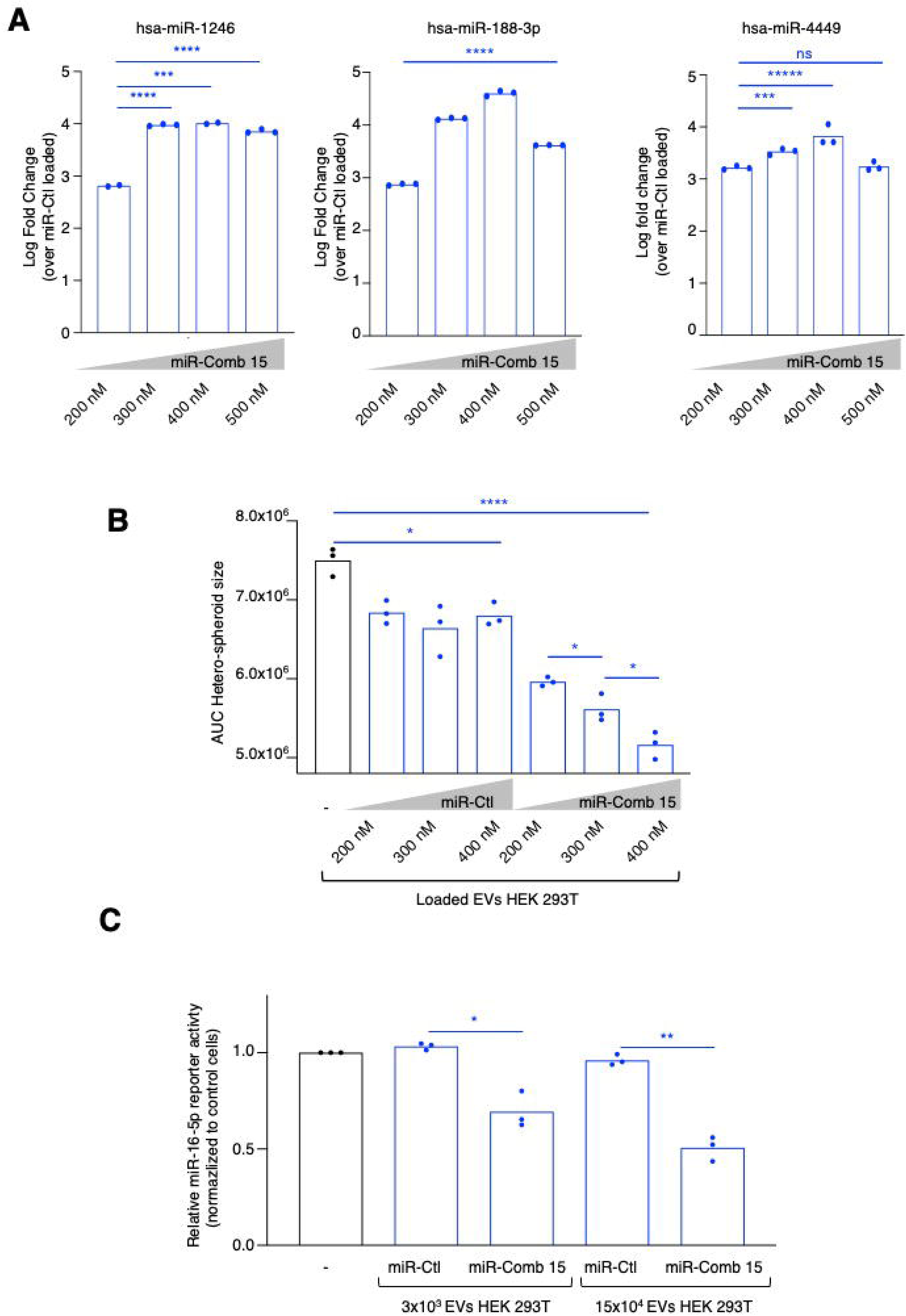
Efficient exogenous loading of miR-Comb 15 enhances the biological activity of HEK 293T-derived EVs. (A) RT-qPCR quantification of selected miR-Comb 15 miRNAs in HEK 293T-derived EVs following pH-gradient loading with increasing concentrations of miR-Comb 15 (200–500 nM). miRNA abundance is shown as log fold change relative to miR-Ctl-loaded EVs. A dose-dependent increase in miRNA loading was observed up to 400 nM, followed by a plateau, suggesting saturation of EVs loading capacity. Additional miRNAs are shown in Figure S6. (B) Functional evaluation of loaded EVs in MDA-MB-231 hetero-spheroid growth assays. HEK 293T-derived EVs loaded with increasing amounts of miR-Comb 15 induced a progressive inhibition of hetero-spheroid growth compared with untreated cells and miR-Ctl-loaded EVs. (C) Functional assessment of miRNA delivery using a miR-16-5p luciferase reporter assay. HEK 293T-derived EVs loaded with miR-Comb 15 significantly reduced reporter activity compared with miR-Ctl-loaded EVs, demonstrating efficient intracellular delivery and target engagement of loaded miRNAs. Data are presented as the mean of three independent biological experiments (n = 3). Statistical significance was assessed using an unpaired two-tailed Student’s *t*-test. ns, not significant; *P* < 0.05, \**P* < 0.01, \*\**P* < 0.001, \*\*\**P* < 0.0001.

RT-qPCR confirmed enrichment of all 15 miRNAs in EV-miR-Comb 15 preparations relative to EV-miR-Ctl (Figure 5A and Figure S6). Under the selected 400 nM loading conditions, Qubit analyses indicated loading efficiencies of 3.29 ± 0.69% and 5.89 ± 1.27% for EV-miR-Comb 15 and EV-miR-Ctl, respectively (Table S2).

We next investigated whether increasing amounts of loaded miR-Comb 15 translated into enhanced biological activity. Hetero-spheroid growth assays were therefore performed using MDA-MB-231 cells treated with EVs loaded with increasing concentrations of miR-Comb 15 (200–400 nM). Interestingly, the loading procedure preserved the intrinsic anti-tumoral activity of HEK 293T-derived EVs, as EV-miR-Ctl preparations continued to inhibit hetero-spheroid growth in vitro (Figure 5B). Notably, exogenous loading with miR-Comb 15 further potentiated this inhibitory effect in a dose-dependent manner, demonstrating that the therapeutic miRNA cargo enhances the endogenous anti-tumoral properties of HEK 293T-derived EVs.

To determine whether EVs-delivered miRNAs remained functionally active following exogenous loading and transfer to recipient cells, we employed a luciferase reporter system containing a miR-16-5p-responsive sequence. As miR-16-5p is one of the fifteen components of miR-Comb 15, this assay provided a means to evaluate the biological activity of the EV-delivered combinatorial miRNA cargo. Treatment with EV-miR-Comb 15 significantly reduced reporter activity compared with EVs-miR-Ctl, and this repression increased proportionally with the amount of EVs-miR-Comb 15 administered to recipient cells (Figure 5C). These findings provide direct functional evidence that EVs-delivered miRNAs remain biologically active following exogenous loading and transfer to recipient cells. Although this assay specifically assessed miR-16-5p activity, the results suggest that the combinatorial miRNA cargo remains functionally active following EV loading and intracellular delivery, as exemplified by the activity of miR-16-5p.

Collectively, these findings establish a robust and scalable strategy for engineering HEK 293T-derived EVs with a therapeutically relevant combinatorial miRNA cargo. Exogenous loading of miR-Comb 15 preserved EVs integrity and miRNA functionality while potentiating the intrinsic anti-tumoral activity of HEK 293T-derived EVs, supporting their further evaluation as RNA delivery platforms for aggressive cancers.

### HEK 293T-derived EVs exhibit the most favorable overall anti-tumoral profile following miR-Comb 15 loading and deliver therapeutic miRNAs to the tumor in vivo

To determine whether these anti-tumoral effects were conserved across additional aggressive cancer models, we extended our analyses to SUM-159PT, and MIA-PaCa-2 cells. Results presented in Figure 6C demonstrate that the anti-tumoral activity of miR-Comb 15-loaded HEK 293T-derived EVs was reproducibly observed across the three aggressive cancer models examined. These data demonstrate that the anti-tumoral activity of miR-Comb 15-loaded HEK 293T-derived EVs is reproducible across multiple aggressive cancer models when compared with untreated cells (black bars)

**Figure 6.**
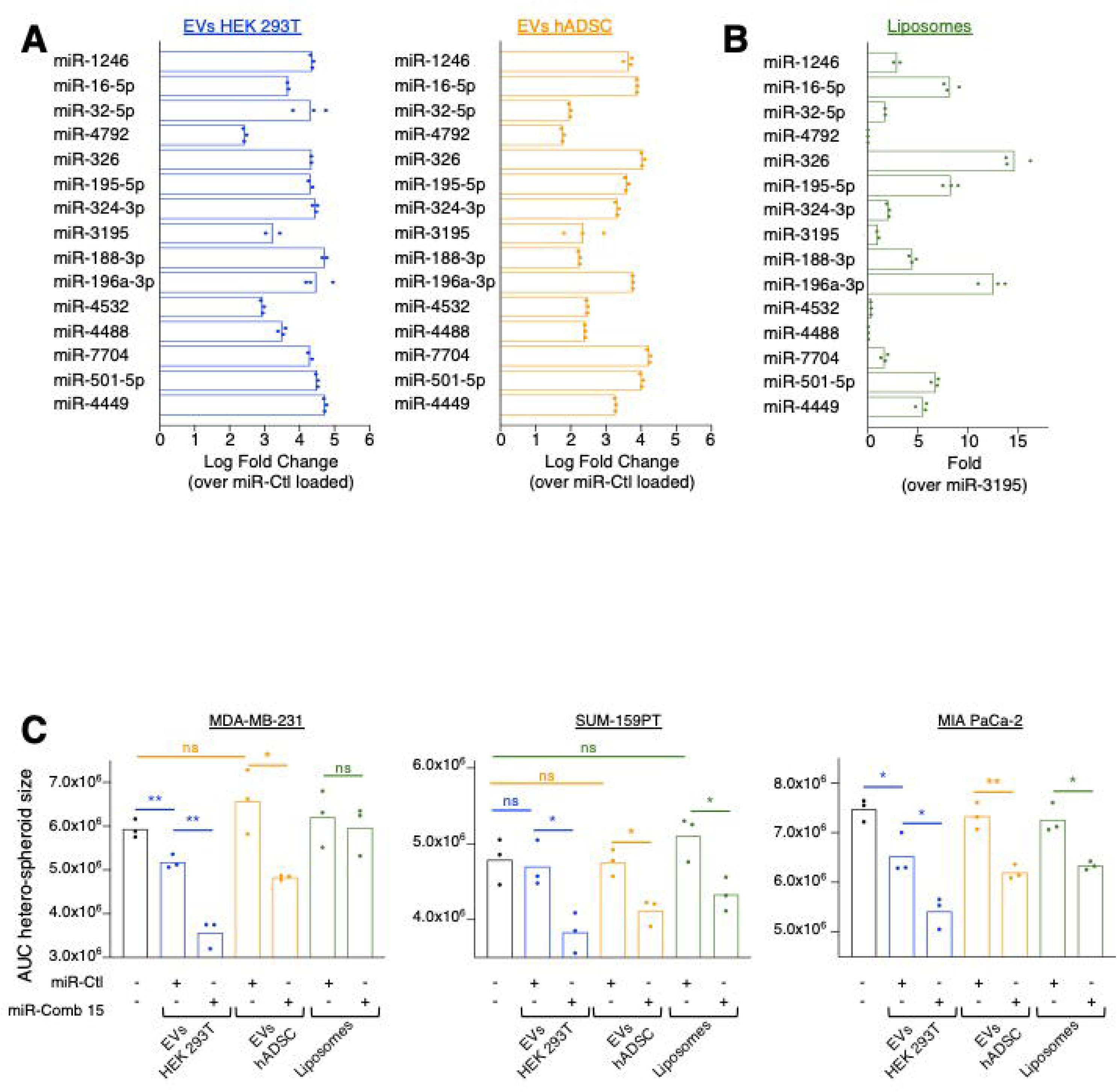
HEK 293T-derived EVs outperform hADSC-derived EVs and liposomes as carriers for miR-Comb 15 delivery in vitro. (A) RT-qPCR quantification of the 15 miRNAs composing miR-Comb 15 after loading into HEK 293T- or hADSC-derived EVs. miRNA levels are expressed as log fold change relative to miR-Ctl-loaded EVs. (B) RT-qPCR quantification of miR-Comb 15 members following loading into liposomes. miRNA levels are expressed relative to miR-3195 (set to 1). Median Ct values are provided in Figure S6. Data are presented as mean from three independent experiments (n = 3). (C) Hetero-spheroid growth assays performed with MDA-MB-231, SUM-159PT, and MIA PaCa-2 cells treated with HEK 293T-derived EVs, hADSC-derived EVs, or liposomes loaded with miR-Ctl or miR-Comb 15. Proliferation was quantified as the area under the curve (AUC). Data are presented as mean from three independent experiments (n = 3). Statistical significance was determined using an unpaired two-tailed Student’s *t*-test (*ns*, not significant; *P* < 0.05; P < 0.01).

In addition, we considered it critical to challenge HEK 293T-derived EVs as miR-Comb-15 carriers against other widely established delivery vehicles. Indeed, we included in our study hADSC-derived EVs, used in therapeutic EVs studies ^46–48^, and liposomes, a synthetic carrier broadly employed for nucleic-acid and drug delivery ^49,50^.

Although these two alternative carriers successfully incorporated the miR-Comb 15 miRNAs (Figure 6A and 6B), RT-qPCR analysis revealed that HEK 293T-derived EVs consistently displayed the highest overall loading efficiency compared to hADSC EVs. Since liposomes lack endogenous miRNA required to calculate a fold change, we compared their miRNA enrichment to the other carriers by assessing each miRNA median Ct (Figure S8). The results in Figure S8 show that liposomes, as well as hADSC EVs, incorporated less miRNA than HEK 293T-derived EVs. Consistently, the carriers loaded with either miR-Ctl or miR-Comb-15 retained indistinguishable size-distribution patterns within each carrier type (Figure S9), indicating that miRNA cargo composition does not alter carriers size properties.

To objectively compare the performance of the different delivery systems, superior efficacy was defined based on a combination of criteria including (i) efficient miRNA loading, (ii) reproducible anti-tumoral activity across multiple aggressive cancer models in vitro, and (iii) the ability to translate these effects into significant tumor growth inhibition in vivo.

Functionally, the in vitro data in Figure 6C show that, among the miR-Ctl loaded conditions, only HEK 293T-derived EVs were able to reduce hetero-spheroids growth in MDA-MB-231 and MIA-PaCa-2 cells but not in SUM-159-PT, indicating that HEK 293T EVs, but not hADSC EVs, exhibit endogenous inhibitory activity in these models. Loading the EVs with miRNA Comb-15 markedly amplified these effects: both HEK 293T EVs and hADSC EVs loaded with miR-Comb-15 significantly inhibited hetero-spheroids growth relative to their miR-Ctl counterparts across all models. In contrast, miR-Comb-15–loaded liposomes induced a clear inhibitory response in SUM-159PT and MIA-PaCa-2 cells but failed to elicit detectable effects in MDA-MB-231 hetero-spheroids growth. Notably, the most consistent anti-tumoral activity across the different cancer models was observed with miR-Comb-15–loaded HEK 293T EVs. Given the reproducible anti-tumoral effects observed across multiple in vitro models, we next sought to determine whether miRNA Comb-15–loaded carriers could elicit comparable effects in vivo. To this end, we used the orthotopic TNBC D3H2LN murine model employed earlier in our present study. One week after tumor cells implantation in athymic nude mice, animals received weekly intravenous injections of either PBS or HEK 293T-derived EVs, hADSC-derived EVs or liposomes loaded with miR-Ctl or the miR-Comb 15. Results presented in Figure 7A show that none of the miR-Ctl loaded carriers significantly inhibited tumor growth compared with the PBS-treated mice. In contrast, both miR-Comb 15–loaded HEK 293T-derived EVs and liposomes significantly inhibited tumor growth; however, HEK 293T-derived EVs displayed a marked advantage, achieving an approximately twofold greater reduction than miR-Comb 15–loaded liposomes relative to PBS-treated controls. Surprisingly, miR-Comb 15 loaded hADSC EVs were not efficient in decreasing tumor growth despite their successful capacity to inhibit hetero-spheroids growth in vitro (Figure 6C).

**Figure 7.**
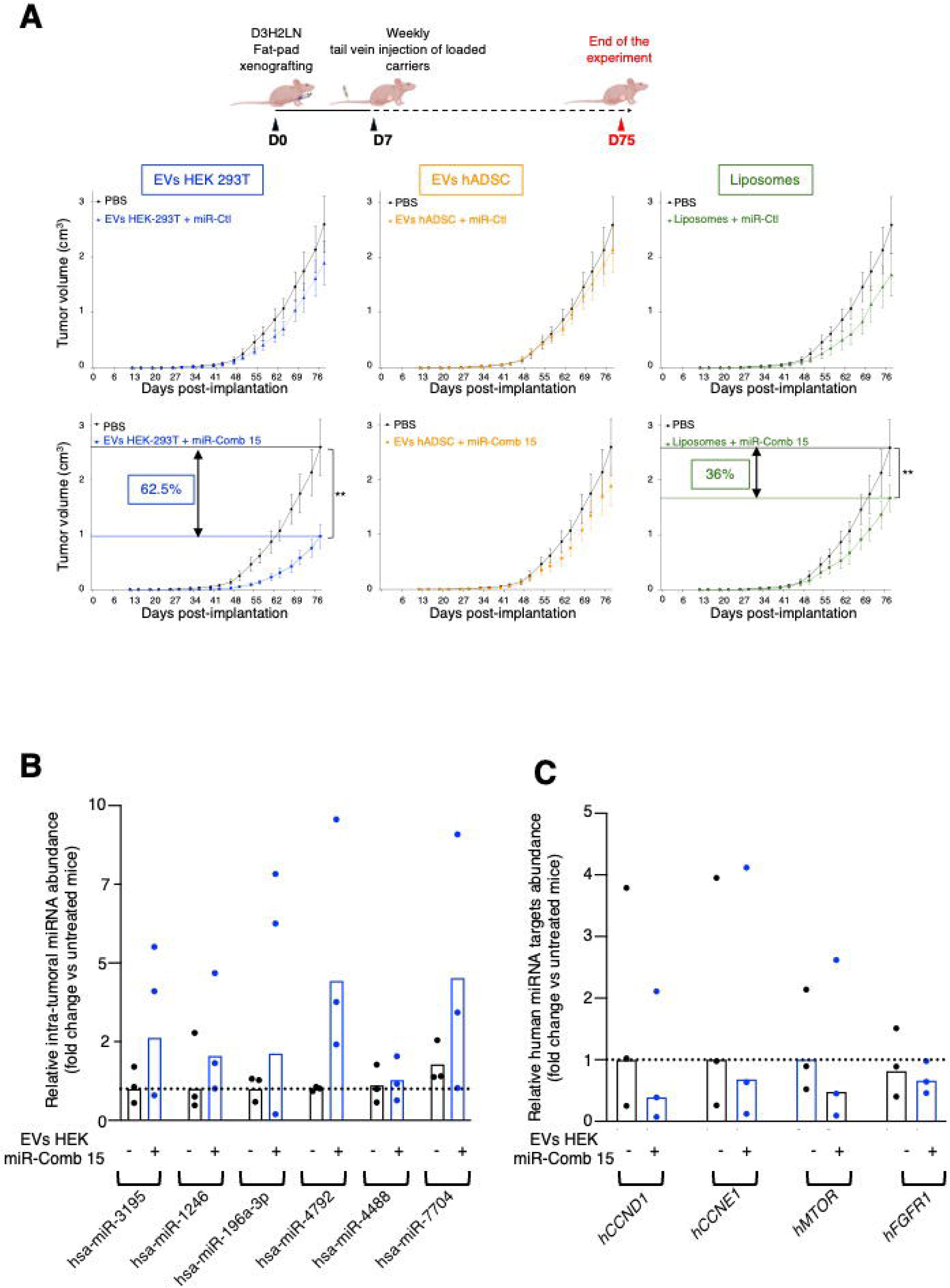
Systemic administration of miR-Comb 15 loaded HEK 293T-derived EVs inhibits tumor growth and promotes intra-tumoral delivery of therapeutic miRNAs in vivo. (A) Experimental design and tumor growth following weekly intravenous administration of HEK 293T-derived EVs, hADSC-derived EVs, or liposomes loaded with miR-Ctl or miR-Comb 15 in an orthotopic D3H2LN TNBC model. Tumor volume was measured by caliper. Data are presented as mean ± SEM (n = 10 mice/group). P < 0.01 (two-way repeated-measures ANOVA). (B) RT-qPCR quantification of selected human-specific miR-Comb 15 members in tumors from mice treated with EV-miR-Comb 15 or control EVs. miRNA levels are expressed relative to control tumors (set to 1). Each dot represents one tumor (n = 3/group). (C) RT-qPCR analysis of representative validated human miR-Comb 15 target genes in the same tumors. Gene expression is expressed relative to control tumors (set to 1). Each dot represents one tumor (n = 3/group).

To investigate whether components of the therapeutic miRNA cargo could be detected within tumors following systemic administration, we quantified, at the end of the experiment, a subset of human-specific miR-Comb 15 members that do not share identical mature sequences with their murine counterparts. Increased levels of multiple human-specific miRNAs were detected in tumors collected from EVs-miR-Comb 15-treated mice compared with PBS-treated controls (Figure 7B), supporting the notion that EV-associated therapeutic miRNAs can reach the tumor microenvironment following intravenous administration. Exploratory analyses revealed trends consistent with the modulation of representative proliferation- and invasion-associated genes in these tumors following miR-Comb 15 treatment (Figure 7C). Although these observations were obtained from a limited number of samples, the overall trend was consistent with our in vitro findings and supports the biological relevance of the selected downstream effectors in vivo.

Importantly, the detection of human-specific members of miR-Comb 15 within tumors following systemic administration provides supportive evidence that therapeutic EV-associated miRNAs reach the tumor microenvironment in vivo. These analyses do not constitute a comprehensive biodistribution assessment. In particular, they do not distinguish whether the detected miRNAs localize within malignant cells or other cellular components of the tumor microenvironment, nor do they provide information regarding the distribution of EVs-miR-Comb 15 across non-tumoral organs. Consequently, direct effects on tumor cells and indirect systemic effects cannot be formally discriminated based on the current dataset. Future studies combining whole-body biodistribution approaches with cell type-specific uptake analyses will therefore be required to fully characterize the in vivo fate of EV-delivered therapeutic miRNAs.

In conclusion, HEK 293T-derived EVs exhibited the most favorable overall performance among the delivery platforms evaluated in this study, combining efficient miRNA loading with reproducible anti-tumoral activity across multiple aggressive cancer models in vitro and robust tumor growth inhibition in vivo.

## Discussion

RNA-based therapies have emerged as a major class of innovative therapeutics, as illustrated by the clinical success of mRNA-based COVID-19 vaccines. In addition to mRNA, multiple therapeutic strategies based on small regulatory RNAs like miRNAs and siRNAs are under active investigation for a wide range of diseases, with several already approved for clinical use^51^. There is a growing interest in RNA-based approaches capable of modulating complex gene networks, a feature particularly relevant for aggressive cancers such as TNBC and pancreatic cancer, which are characterized by marked molecular heterogeneity and simultaneous activation of multiple oncogenic pathways ^52,53^. In this context, EVs have gained considerable attention as natural carriers for therapeutic RNAs owing to their ability to protect RNA cargo from degradation, mediate functional intercellular transfer, and potentially offer improved biocompatibility compared with synthetic delivery systems^54^.

We previously identified NFAT3 as a potent anti-tumoral transcription factor ^26,27^ in luminal breast cancer cells and demonstrated that EVs derived from NFAT3-expressing cells inhibit in vitro invasion and proliferation across multiple cancer models^28^. Furthermore, in vivo, NFAT3 also inhibits tumor growth and metastatic dissemination in a mouse TNBC model ^28^. Notably, these effects were obtained despite the absence of detectable NFAT3 mRNA or protein transfer to recipient cells ^28^. These findings strongly suggested that NFAT3 may contribute to its tumor-suppressive effects by shaping the anti-tumoral cargo composition of EVs rather than through direct transfer of NFAT3 itself. Consistent with this hypothesis, the present study identified a 15-miRNA signature (miR-Comb 15) that functionally recapitulates the anti-tumoral activity of NFAT3-regulated EVs. Moreover, our new data demonstrate that NFAT3 depletion alters the expression of several members of this signature (Figure S1), providing additional evidence that NFAT3 contributes to the establishment of an anti-tumoral EV-associated miRNA program. However, the molecular mechanisms linking NFAT3 activity to miR-Comb1 5 expression and EVs cargo composition remain to be elucidated.

Interestingly, while several miRNAs displayed autonomous anti-proliferative activity, none of the anti-migratory miRNAs was individually sufficient to inhibit cell invasion. These observations indicate that the full inhibitory effect relies on the coordinated action of the entire set of miRNAs. Beyond functional assays, direct transfection of miR-Comb 15 induced marked morphological changes, including increased cell area in breast and pancreatic cancer cells. Such alterations in cell size have been associated with reduced proliferative and invasive capacities^43,44^, supporting the notion that miR-Comb15 impacts fundamental cellular properties and may contribute to a broader reprogramming of cellular behavior. Our findings provide experimental support for the concept that coordinated modulation of multiple miRNAs converges on the regulation of key proliferative and invasive programs rather than on the inhibition of a single dominant target. By integrating experimentally validated miRNA-target interactions, functional enrichment analyses, and experimental validation of representative downstream genes, our study provides a mechanistic framework supporting the network-modulating properties of combinatorial miRNA therapies. Exploratory analyses performed in tumors from EVs-miR-Comb 15-treated mice revealed trends consistent with the molecular changes observed in vitro, further supporting the potential biological relevance of these downstream effectors in vivo, although confirmation in larger studies will be required.

The cellular targets responsible for the anti-tumoral effects mediated by miR-Comb 15 remain to be fully elucidated. Our previous work demonstrated that macrophages were required for the anti-proliferative activity, but not the anti-invasive activity, of NFAT3-regulated EVs in hetero-spheroid models^28^. Consistent with these findings, miR-Comb15 inhibited tumor cell invasion in conventional invasion assays performed in the absence of macrophages, supporting a direct tumor cell-intrinsic mechanism. Nevertheless, the current study does not allow discrimination between direct actions of miR-Comb 15 on macrophages and indirect effects arising from miR-Comb 15-induced alterations in tumor cell behavior and tumor–microenvironment interactions. Further investigations will therefore be necessary to dissect the respective contributions of tumor cell-intrinsic and microenvironment-dependent mechanisms to the therapeutic activity of miR-Comb 15. Importantly, because the in vivo experiments were performed in athymic mice retaining innate immune populations, including macrophages, a contribution of tumor–microenvironment interactions to the overall therapeutic response cannot be excluded.

Because our previous work demonstrated that NFAT3-regulated EVs failed to inhibit tumor cell proliferation in conventional 2D monocultures while retaining anti-tumoral activity in more physiologically relevant settings and in vivo^28^, hetero-spheroid models were deliberately employed in the present study. By incorporating tumor-stroma interactions, these models better recapitulate key aspects of the tumor microenvironment and therefore provide a biologically relevant framework to assess the functional consequences of miR-Comb 15 treatment. Furthermore, the anti-tumoral effects observed in hetero-spheroid assays were subsequently extended in an orthotopic TNBC model, supporting the translational relevance of this experimental strategy.

The functional relevance of EVs-associated miRNAs has been debated, with reports suggesting that miRNAs are minor EVs constituents^41,55,56^ and inefficiently delivered to recipient cells^41,55,56^. Our results refine this view by showing that network complexity is a critical determinant of biological impact. Although individual miRNAs may be present at low stoichiometry^57^, their coordinated action can be sufficient to induce robust phenotypic effects. Importantly, our study provides functional evidence consistent with the transfer of biologically active miRNAs from EVs to recipient cells. Indeed, transfection of recipient cells with specific antagomirs abrogated the phenotypic effects induced by NFAT3-regulated EVs, supporting the notion that EV-associated miRNAs contribute to the observed biological responses. Nevertheless, direct visualization of EVs uptake, precise intracellular localization of EV-delivered miRNAs, and formal demonstration of RISC engagement were not addressed in the present study and will require dedicated future investigations.

From a translational perspective, the use of cancer cell-derived EVs raises safety and regulatory concerns^58^. Although T47D-derived EVs enabled the identification of the anti-tumoral miRNA signature described in this study, their malignant origin raises important translational concerns. Tumor-derived EVs may contain additional bioactive cargo, including oncogenic proteins, nucleic acids, and signaling molecules capable of promoting tumor progression ^59^, metastatic niche formation^60^, immune modulation, or therapy resistance^61^. Therefore, despite their utility as a discovery platform, the direct therapeutic application of tumor-derived EVs remains questionable.

We therefore selected HEK 293T-derived EVs as an alternative platform more compatible with therapeutic development. HEK 293T cells are characterized by robust growth properties, high EVs production yields, ease of genetic manipulation, and extensive use in EVs engineering studies.

To further enhance the therapeutic potential of HEK 293T EVs while avoiding genetic manipulation of producer cells, we employed a pH-gradient-based loading strategy^40^ that enabled efficient, saturable incorporation of miR-Comb 15 without altering EVs size or intrinsic bioactivity. This approach proved compatible with the simultaneous loading of multiple miRNAs, a prerequisite for combinatorial RNA therapies, and provided a practical strategy for controlled EVs cargo engineering. In contrast, hADSC-derived EVs and liposomes loaded with miR-Comb 15 displayed only modest efficacy, highlighting EVs origin as a critical determinant of therapeutic outcome. Collectively, these findings establish HEK 293T-derived EVs as a scalable and engineerable platform for the development of combinatorial RNA therapeutics.

Interestingly, the anti-tumoral activity of HEK 293T-derived EVs did not increase proportionally with EVs dose, as lower EVs concentrations produced the strongest biological effects in certain experimental settings. Although the mechanisms responsible for this non-linear dose-response relationship were not investigated in the present study, previous reports indicate that EVs uptake and cargo delivery rely on complex and often inefficient endocytic processes that may display non-linear behaviors depending on EVs concentration, recipient cell type and intracellular trafficking pathways ^62–64^. Additional studies will therefore be required to elucidate the mechanisms underlying this phenomenon.

Because the same miR-Comb15 cargo was delivered using HEK 293T-derived EVs, hADSC-derived EVs, and synthetic liposomes, the differences observed in vivo are likely attributable to intrinsic characteristics of the delivery vehicles rather than to the therapeutic cargo itself. EVs origin and composition may influence membrane properties, surface-associated proteins, circulation stability, biodistribution, cellular tropism, and interactions with the tumor microenvironment, thereby shaping the biological fate and therapeutic efficacy of administered vesicles. Further studies will be required to define the mechanisms underlying these differences and to identify the EVs sources best suited for therapeutic applications.

Future studies addressing large-scale manufacturing, biodistribution, long-term safety, and characterization of engineered EVs preparations in accordance with MISEV2023 recommendations will be required before clinical translation can be envisaged.

An additional consideration concerns the intrinsic biological activity previously reported for HEK 293T-derived EVs. Importantly, EV-miR-Ctl failed to significantly inhibit tumor growth in vivo, whereas EVs-miR-Comb 15 induced robust anti-tumoral effects. Because both preparations originated from the same HEK 293T-derived EVs platform and underwent identical loading procedures, these observations strongly support an additional therapeutic contribution of the miRNA cargo beyond the intrinsic properties of the EVs carrier.

In summary, this work identifies a transcription factor-driven EV-miRNA network that underlies EV-mediated tumor suppression in aggressive cancers. Our findings support a model in which NFAT3 shapes an anti-tumoral EVs program through the regulation of miR-Comb 15, whose coordinated activity is required to inhibit tumor cell proliferation and invasion. These observations further emphasize the importance of combinatorial miRNA networks in controlling complex oncogenic phenotypes.

From a translational perspective, our study demonstrates that this anti-tumoral program can be rationally developed into HEK 293T-derived EVs, a scalable and engineerable delivery platform compatible with the development of EV-based RNA therapeutics. The successful loading of miR-Comb 15 while preserving EVs integrity and biological activity further supports the feasibility of this approach.

Finally, these findings have important therapeutic implications. By enabling the simultaneous modulation of multiple oncogenic pathways, EV-mediated delivery of combinatorial miRNA signatures may represent a promising alternative to conventional single-target strategies for the treatment of aggressive malignancies such as TNBC and pancreatic cancer.

## Materials and methods

### Cell lines and culture conditions

Human and murine cell lines were used in this study. MDA-MB-231 (American Type Culture collection) and SUM-159-PT (provided by Alex Toker, Harvard Medical School) are human TNBC cell lines derived from metastatic and inflammatory breast carcinomas, respectively. MIA-PaCa-2 is a human pancreatic ductal adenocarcinoma cell line (American Type Culture collection). RAW 264.7 is a murine macrophage cell line (American Type Culture collection). HEK 293T cells are human embryonic kidney-derived cells expressing the SV40 large T antigen (American Type Culture collection). Two independent shRNA constructs targeting NFAT3 were used to generate stable T47D knockdown cell lines previously described^28^, these cells were used for RT-qPCR validation of NFAT3-dependent miRNAs shown in Figure S1. MDA-MB-231 and SUM-159-PT cells were cultured in DMEM containing 1 g/L glucose, L-glutamine, HEPES, and sodium pyruvate (Gibco, #22320022), supplemented with 10% fetal bovine serum (FBS) (Dustcher), 100 U/mL penicillin/streptomycin (Thermo Fisher Scientific, #15140122). HEK 293T, MIA-PaCa-2, and RAW 264.7 cells were maintained in high-glucose DMEM (4.5 g/L; Gibco, #11960044) supplemented with 10% FBS, penicillin/streptomycin, and L-glutamine. Cells were maintained at 37°C in a humidified atmosphere with 5% CO₂. Normocin™ antimicrobial reagent (InvivoGen, #ant-nr-2) was used during routine maintenance but omitted during experiments. All cell lines were regularly tested for mycoplasma contamination.

### Extracellular vesicles production and isolation and liposomes

Different EVs sources were employed in this study according to their specific experimental purposes. T47D-derived EVs served as a discovery platform for the identification of the NFAT3-regulated anti-tumoral miRNA signature. hADSC-derived EVs were included as a translational comparator because of their widespread use in therapeutic EVs studies. Finally, HEK 293T-derived EVs were selected as the principal therapeutic platform owing to their favorable characteristics for EVs engineering, including high production yield and compatibility with large-scale manufacturing approaches.

T-47D-derived EVs and HEK 293T-derived EVs were isolated by differential centrifugation and ultracentrifugation as previously described^28^. Briefly, cells were cultured to ∼70% confluence and incubated for 48 h in EVs-depleted medium. Conditioned medium was sequentially centrifuged at 300 g, 2,000 g, and 10,000 g to remove cells and debris, followed by ultracentrifugation at 120,000 g for 90 min at 4°C. EVs pellets were washed twice in PBS and re-pelleted under identical conditions. Final EVs preparations were resuspended in PBS, aliquoted, and stored at -80°C. Particle size distribution and concentration were assessed by nanoparticle tracking analysis (NTA) using a ZetaView instrument (Particle Metrix). EVs dosing was normalized according to particle number in all in vitro and in vivo experiments, thereby minimizing variability associated with EVs quantification and ensuring comparability between treatment groups. EVs characterization was performed in accordance with the recommendations of the MISEV2023 guidelines, using nanoparticle tracking analysis to assess particle concentration and size distribution. Additional characterization of T47D-derived EVs, including the evaluation of established EV-associated protein markers, has been previously reported by our group^28^. EVs preparations were used within one freeze–thaw cycle.

Human adipose-derived stem cell (hADSC) EVs were provided by EVerZom and produced using a proprietary large-scale 3D culture and tangential flow filtration process^65^. Plain liposomes (Cellsome®) were purchased (Encapsula Nanoscience, #CEP-500).

### miRNA loading into extracellular vesicles

For miRNA loading experiments, HEK 293T-derived EVs were loaded with a combination of 15 miRNAs using a pH-gradient-based protocol adapted from Jeyaram et al.^40^. Briefly, EVs (9×10^6^ particles) were transiently dehydrated in 100% ethanol at room temperature using a Eppendorf® vacuum Concentrator Plus machine (SpeedVac) for 3 h. Following dehydration, EVs were resuspended in an acidic citric acid buffer (pH 2.5) and incubated for 1 h under gentle agitation at room temperature. Rehydrated EVs were subsequently dialyzed overnight against HBS buffer (pH 7.0) using Float-A-Lyser G2 devices (MWCO 8–10 kDa; Spectrum Laboratories) to restore physiological pH. After dialysis, EVs were concentrated to a final volume of 40 µL using 10 kDa Amicon Ultra 0.5 mL centrifugal filters (Merck).

For the loading procedure, 40 µL of concentrated EVs were incubated for 2 h at room temperature in a final volume of 200 µL of HBS containing miRIDIAN miRNA mimics (Horizon/Dharmacon; sequences listed in Table S1). miRNA concentrations ranging from 200 to 500 nM were evaluated using either miR-Comb 15 or the control miRNA combination (miRNA Ctl). Following incubation, loaded EVs were purified to remove unincorporated miRNAs by successive centrifugation steps using 10 kDa Amicon Ultra 0.5 mL devices, resuspended in PBS, and stored at −80°C until use. Loading efficiency was assessed by RT-qPCR, and saturation kinetics were determined by comparing EVs-associated miRNA levels obtained across the different miRNA concentrations tested.

### RNA extraction and RT-qPCR

#### - Quantification of miRNAs

Total RNA was isolated from cultured cells using the miRNeasy Mini Kit (QIAGEN, #217004). Total RNA from EVs preparations was isolated using the miRNeasy Micro Kit (QIAGEN, #217084). For xenograft tumor analyses, frozen tumor tissues were homogenized in QIAzol Lysis Reagent (QIAGEN), and total RNA, including small RNAs, was extracted using the miRNeasy Micro Kit according to the manufacturer’s instructions. Reverse transcription was performed using the miRCURY LNA™ RT Kit (QIAGEN, #339340), and quantitative PCR was carried out using the miRCURY LNA™ SYBR Green PCR Kit (QIAGEN, #339346) on a 7500 Fast Real-Time PCR System (Applied Biosystems). Relative miRNA expression was calculated using the 2×-ΔΔCt method after normalization to SNORD48. Human-specific LNA assays were used for xenograft tumor analyses to selectively detect therapeutic miRNAs without cross-reactivity with murine homologs.

#### Quantification of human mRNAs

Total RNA extracted from cultured cells or xenograft tumors was reverse transcribed using the iScript™ cDNA Synthesis Kit (#1708891, Bio-Rad) according to the manufacturer’s instructions. Quantitative PCR was performed using human-specific primer assays (AnyGenes, Paris, France) on a 7500 Fast Real-Time PCR System (Applied Biosystems). Relative gene expression was calculated using the 2×−ΔΔCt method after normalization to the geometric mean of HPRT1, B2M, and GAPDH.

### miRNA loading efficiency

For the EVs preparations generated under the 400 nM loading conditions used throughout this study, miRNA loading efficiency was estimated using the Qubit microRNA Assay Kit (Thermo Fisher Scientific). Following the loading procedure and purification steps, total RNA was extracted from engineered EVs preparations using the same protocol described above. The amount of RNA recovered from EVs preparations was then compared with the initial amount of input miRNA used during loading, and loading efficiency was calculated Loading efficiency was calculated according to Equation (1).

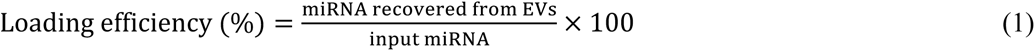

Measurements were performed with technical replicates from two independent loading experiments.

### Scratch wound healing assay

MDA-MB-231 cells were seeded in 96-well plates and transfected with individual miRNA antagomirs or a non-targeting control antagomir (sequences in Table S1) using DharmaFECT 1 reagent (Horizon/Dharmacon, #T-2001-03) according to the manufacturer’s instructions. Twenty-four hours after transfection, a scratch wound was generated in each well using a standardized wound-making tool. Cells were then treated with either extracellular vesicles in appropriate medium or medium as a control condition. Wound closure was monitored by live-cell imaging using the IncuCyte S3 system (Essen Bioscience), with images acquired every hour.

### 3D hetero-spheroids growth assay

Cancer cells were transfected or not with either individual mimics miRNA or mimics miRNA combinations or individual antagomirs (sequences in Table S1) as indicated in the figures ‘legends. The following day cells were co-cultured with RAW 264.7 macrophages at a 2:1 ratio to generate hetero-spheroids. Cell mixtures were embedded in 3% Matrigel and seeded in ultra-low attachment plates. After hetero-spheroids formation, they were either untreated or incubated with EVs or liposomes, and growth was monitored for 96 h using live-cell imaging. Hetero-spheroids area was quantified over time using the IncuCyte S3 system (Essen Bioscience) every 2 hours for 4 days. AUC of hetero-spheroids growth was calculated with the GraphPad Prism software for each tested condition for a total time of 4 days, as previously described^28^.

### In vitro invasion assay

Cells transfected with miRNAs were seeded into Matrigel-coated transwell inserts. NIH/3T3-conditioned medium was used as a chemoattractant. After 6 h of incubation, invading cells were fixed, stained, and quantified by microscopy. Within each experiment, each condition was tested in triplicates and the number of cells in the transwell whole field was counted to evaluate the cells’ ability to invade the Matrigel. Thus, the invasion index was calculated as the proportion of the number of invading cells in treated wells compared to the number of invading cells in the control wells (-) arbitrarily set to 1. Invasion assays were performed using Matrigel-coated transwell inserts, without macrophage co-culture.

### Immunofluorescence staining

Cells were seeded on glass coverslips in 12-well plates and cultured in their respective medium supplemented with 10% FBS. Twenty-four hours after seeding, cells were transfected with either a non-targeting control miRNA or mimic miRNAs using DharmaFECT 1 reagent (Horizon/Dharmacon, #T-2001-03) according to the manufacturer’s instructions. Transfected cells were maintained under standard culture conditions and processed for immunofluorescence staining 48 h post-transfection. Cells were fixed with 4% paraformaldehyde, permeabilized with 0.1% Triton X-100, and quenched with 50 mM NH₄Cl. After blocking with PBS-Tween containing 3% BSA, cells were incubated with Alexa Fluor 488 phalloidin (Cell Signaling Technology, #8878) for 1 h, followed by DAPI nuclear staining (Cell Signaling Technology, #62248). Coverslips were mounted using FluoroMount-G (SouthernBiotech, #0100-01) and imaged with a ZEISS LSM800 confocal laser microscope equipped with a Plan-Apochromat 40×/1.3 NA oil immersion objective using ZEN software.

### In vivo tumor studies

All animal experiments were conducted in accordance with institutional guidelines and were approved by the relevant ethical committee under protocol number #21058. Female mice were used because breast tumor cells were implanted into the mammary fat pad, an orthotopic site that provides a physiologically relevant microenvironment for breast cancer growth. In addition, breast cancer predominantly affects women, further supporting the use of female animals. Indeed, six-week-old Hsd: Athymic Nude-Foxn1nu female mice were purchased from Charles Rivers company and used to establish an orthotopic TNBC model. Briefly, D3H2LN cells were implanted orthotopically into the mammary fat pad as previously described^28^. One week after tumor cell implantation, mice were randomly assigned to treatment groups and received weekly intravenous injections of either PBS or vesicular carriers loaded with control miRNA (miR-Ctl) or miR-Comb 15. The carriers tested included HEK 293T-derived EVs, hADSC-derived EVs, and liposomes as indicated in the figures’ legends. Tumor growth was monitored over time by caliper measurements, and tumor volumes were calculated using standard formulas. At the end of the experiments, mice were sacrificed by cervical dislocation. Some tumors were kept frozen at -80°C.

### RNA sequencing

EVs from T47D shCtl and T47D shNFAT3 cells were isolated as described above. Total RNA was extracted using the Rneasy Plus Micro Kit (QIAGEN, #74034) and assessed for quality prior to library preparation and sequencing on an Illumina NextSeq 550 platform. Bioinformatic analyses were performed by GenoSplice (http://www.genosplice.com/) using CAP-miRSeq^66^.

### Image and statistical analysis

Image analysis was performed using ImageJ. Statistical analyses were conducted using GraphPad Prism 10 software (GraphPad Software, Inc., San Diego, CA, USA). P value designations were as follows: ns = not significant or p ≥ 0.05, *p < 0.05, **p < 0.005, ***p < 0.001, and ****p < 0.0001.

### Study Approval

All animal experiments were performed in accordance with animal care ethics approval and guidelines of the Université Paris Cité Ethic Committee. All procedures were approved under the Protocol #21058 (Jauliac Laboratory) by the Université Paris Cité Ethic Committee.

## Data availability statement

miRNA sequencing data associated with this paper (both processed counts and raw data) are publicly available via GEO: GSE318086

## Acknowledgments

We thank the members of Dr. Lehmann-Che and Dr. Jauliac laboratory for insightful discussions. We thank members of the technological platform of IRSL for technical advice. We thank EVerZom for hADSC EVs and access to their platform to evaluate EVs production quality and concentration.

This work was supported by the SATT INNOV IDF (ERGANEO), Ligue Nationale contre le Cancer, Gefluc Paris, INSERM, and Université Paris Cité. We thank the french ministry of scientific research that granted Guénolé Tossou a PhD scholarship.

All the authors have read and approved the final manuscript for publication.

## Authors contributions

Conceptualization, Sébastien Jauliac, Guénolé Tossou.; Methodology, Sébastien Jauliac, Guénolé Tossou., Nadia Ourari, Maëlle Ralu, Alexandre Montanede., Frédéric Guaddachi., Babette Beher, Manon Paul, Emilie Ouanounou., Stéphane Brunet; Writing – review & editing, Sébastien Jauliac, Guénolé Tossou, Stéphane Brunet, Morgane le Bras, Jacqueline Lehmann-Che, Maëlle Ralu; Supervision, Sébastien Jauliac; Funding acquisition, Sébastien Jauliac

## Consent for publication

