## Supplementary material for "A combinatorial EVs-miRNA signature mediates the anti-tumoral activity of NFAT3-regulated extracellular vesicles in aggressive cancers": supplemmental material

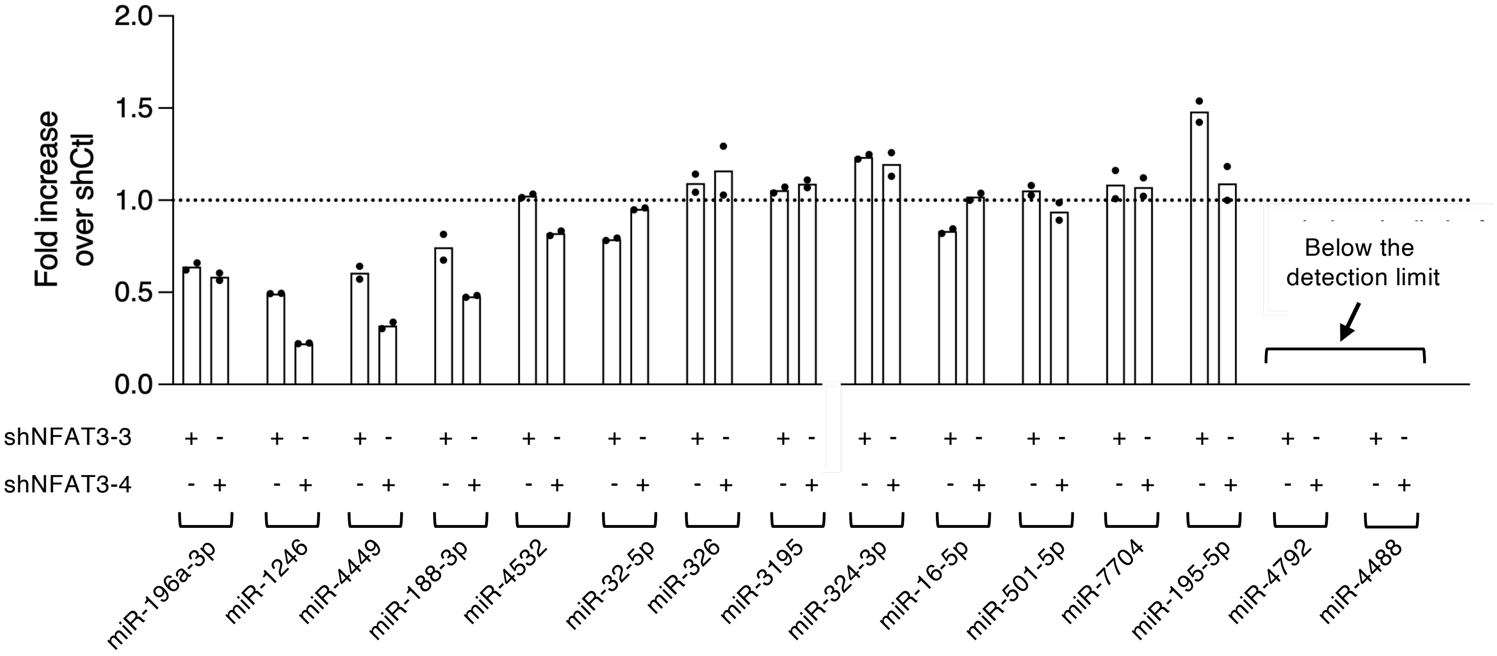


**Figure S1. NFAT3-dependent modulation of miR-Comb15 expression in T47D cells.** RT-qPCR analysis of the 15 miRNAs composing miR-Comb15 in two independent NFAT3 knockdown T47D cell lines (shNFAT3-3 and shNFAT3-4). Data are presented as fold change relative to shCtl cells (set to 1; dotted line) and were calculated using the $2^{-\Delta\Delta Ct}$ method. Data represent two independent experiments (n = 2). miR-4792 and miR-4488 were below the detection limit.

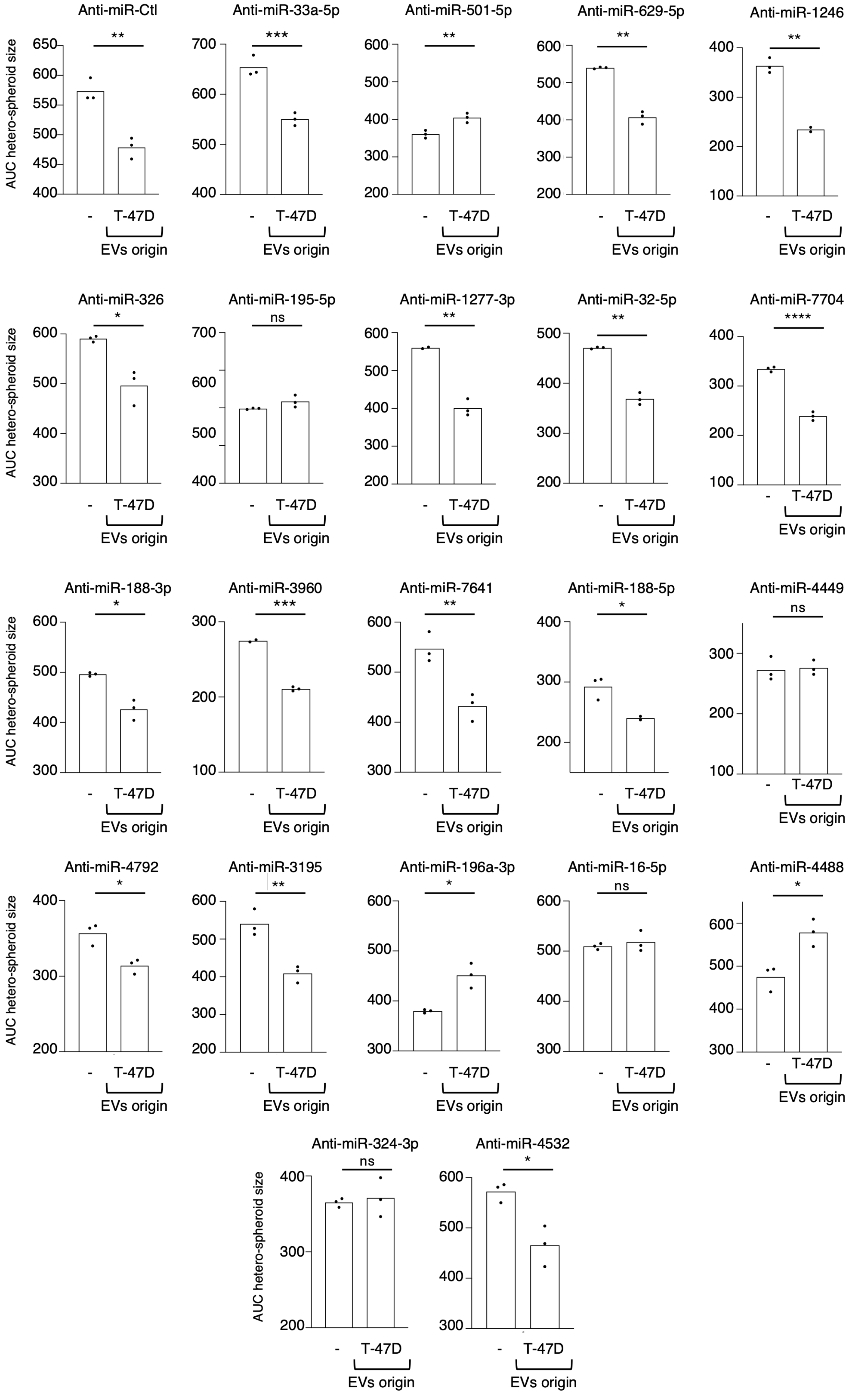
Anti-miR-Ctl
**
Anti-miR-33a-5p
***
Anti-miR-501-5p
**
Anti-miR-629-5p
**
Anti-miR-1246
**
Anti-miR-326
*
Anti-miR-195-5p
ns
Anti-miR-1277-3p
**
Anti-miR-32-5p
**
Anti-miR-7704
****
Anti-miR-188-3p
*
Anti-miR-3960
***
Anti-miR-7641
**
Anti-miR-188-5p
*
Anti-miR-4449
ns
Anti-miR-4792
*
Anti-miR-3195
**
Anti-miR-196a-3p
*
Anti-miR-16-5p
ns
Anti-miR-4488
*
Anti-miR-324-3p
ns
Anti-miR-4532
*
AUC hetero-spheroid size
-
T-47D
EVs origin

**Figure S2. Contribution of individual miRNAs to the anti-tumoral activity of T47D-derived EVs.**

Hetero-spheroid growth assays were performed using MDA-MB-231 cells treated with EVs isolated from T-47D cells transfected with individual antagomir (anti-miRNAs) targeting members of the miR-Comb 15 signature or with a negative control antagomir (Anti-miR-Ctl). Proliferation was quantified as the area under the curve (AUC) of hetero-spheroid growth. Data are presented as the mean of three independent biological experiments (n = 3). Statistical significance was assessed using an unpaired two-tailed Student's *t*-test. ns, not significant; $P < 0.05$; *$P < 0.01$; **$P < 0.001$; ***$P < 0.0001$.

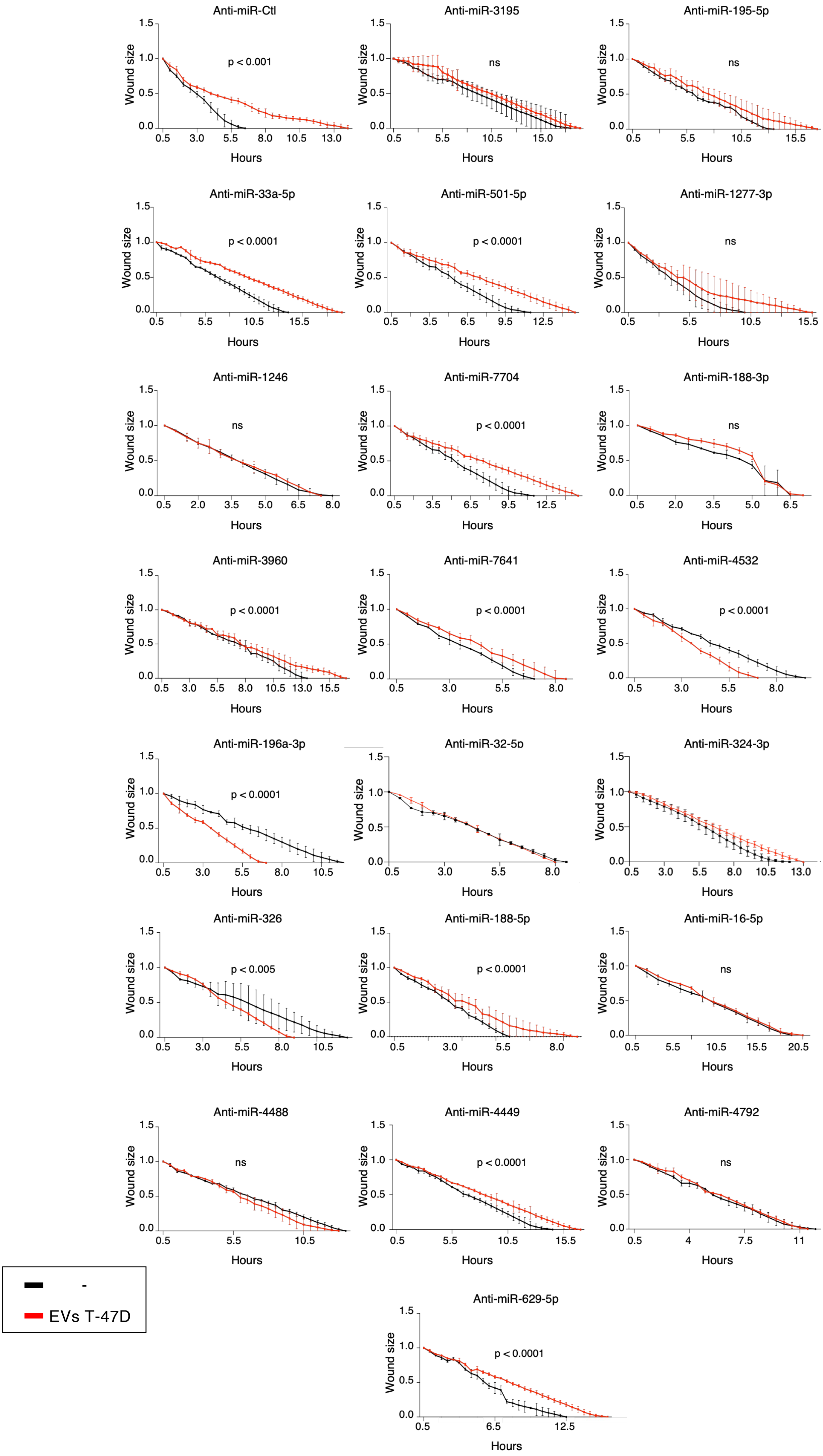
Anti-miR-Ctl
p < 0.001
Anti-miR-3195
ns
Anti-miR-195-5p
ns
Anti-miR-33a-5p
p < 0.0001
Anti-miR-501-5p
p < 0.0001
Anti-miR-1277-3p
ns
Anti-miR-1246
ns
Anti-miR-7704
p < 0.0001
Anti-miR-188-3p
ns
Anti-miR-3960
p < 0.0001
Anti-miR-7641
p < 0.0001
Anti-miR-4532
p < 0.0001
Anti-miR-196a-3p
p < 0.0001
Anti-miR-32-5p
Anti-miR-324-3p
Anti-miR-326
p < 0.005
Anti-miR-188-5p
p < 0.0001
Anti-miR-16-5p
ns
Anti-miR-4488
ns
Anti-miR-4449
p < 0.0001
Anti-miR-4792
ns
Anti-miR-629-5p
p < 0.0001
Wound size
Hours
-
EVs T-47D

**Figure S3. Contribution of individual miRNAs to the anti-migratory activity of T47D-derived EVs.**

Wound-healing assays were performed using MDA-MB-231 cells treated with EVs isolated from T-47D cells transfected with individual antagomir (anti-miRNA) targeting members of the miR-Comb 15 signature or with a negative control antagomir (Anti-miR-Ctl). Wound closure was monitored over time using live-cell imaging and quantified as relative wound size. Black curves correspond to untreated cells and red curves to cells treated with T-47D-derived EVs. Data are presented as mean ± SEM from three independent biological experiments ($n = 3$). Statistical significance was assessed using two-way repeated-measures ANOVA; ns, not significant; $P < 0.05$; $P < 0.01$; $P < 0.001$; $P < 0.0001$.

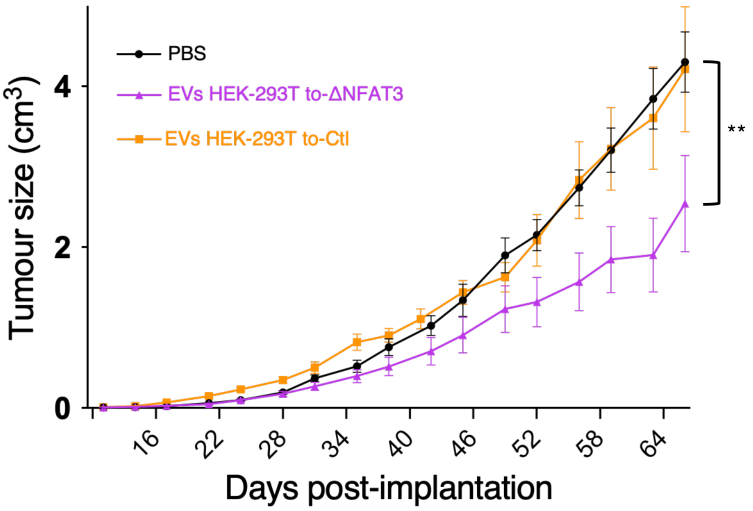


**Figure S4. Anti-tumoral activity of EVs derived from NFAT3-expressing cells in vivo.**

Tumor growth was monitored by caliper measurements following orthotopic implantation of D3H2LN cells into the mammary fat pad of mice. Animals received weekly tail-vein injections of PBS, EVs derived from stable HEK-293T (EVs HEK293T to-Ctl), or EVs derived from stable HEK 293T cells expressing constitutively active NFAT3 (EVs HEK293T to-ΔNFAT3), as indicated. Tumor volumes were measured over time and are presented as mean ± SEM. Statistical significance was assessed using two-way repeated-measures ANOVA. $^{**}P < 0.01$.

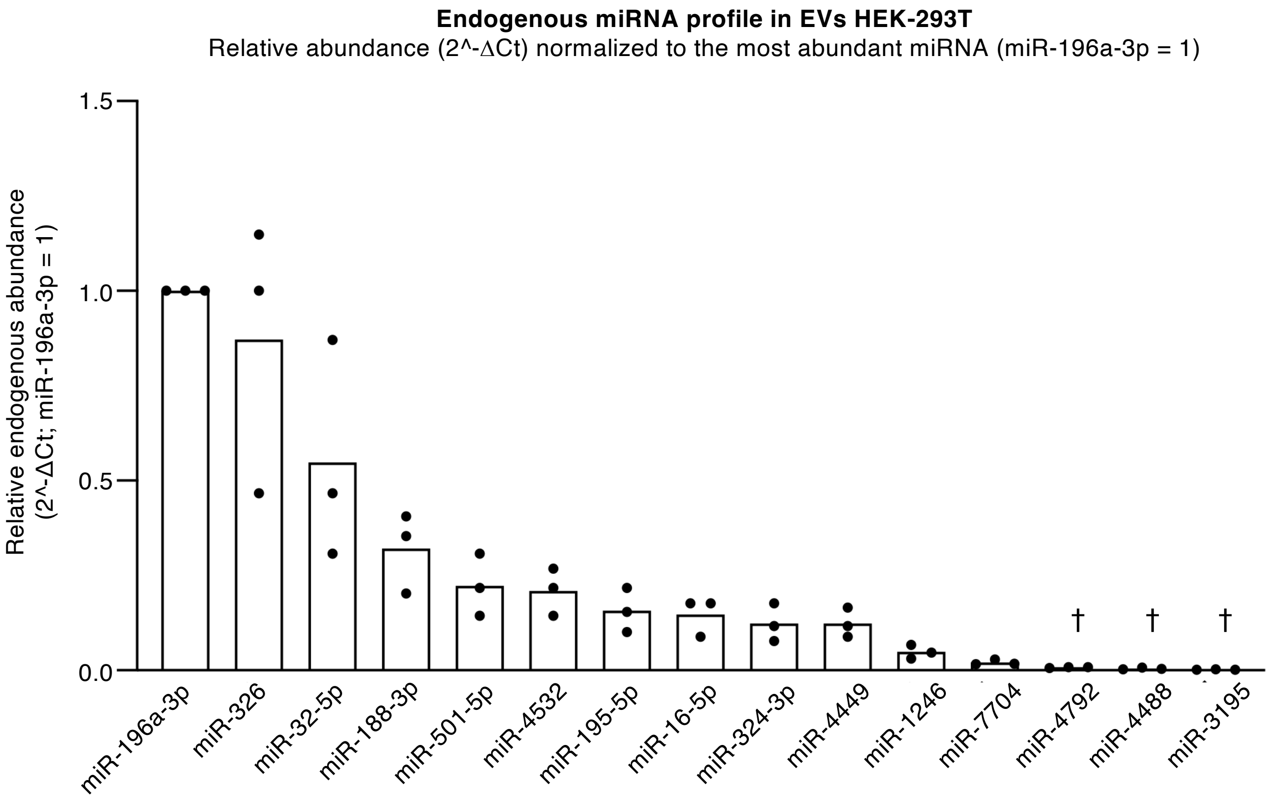


† miRNAs displaying Ct ≥ 35 in at least one independent EV preparation should be interpreted cautiously due to their proximity to the assay detection limit.

**Figure S5**. Endogenous abundance of miR-Comb 15 members in naïve HEK 293T-derived EVs. RT-qPCR analyses were performed to quantify the endogenous levels of the 15 miRNAs constituting miR-Comb15. Relative abundance was calculated using the 2^-ΔCt method and normalized to the most abundant miRNA detected (miR-196a-3p = 1). Data are presented as mean ± SEM from three independent biological experiments (n = 3).

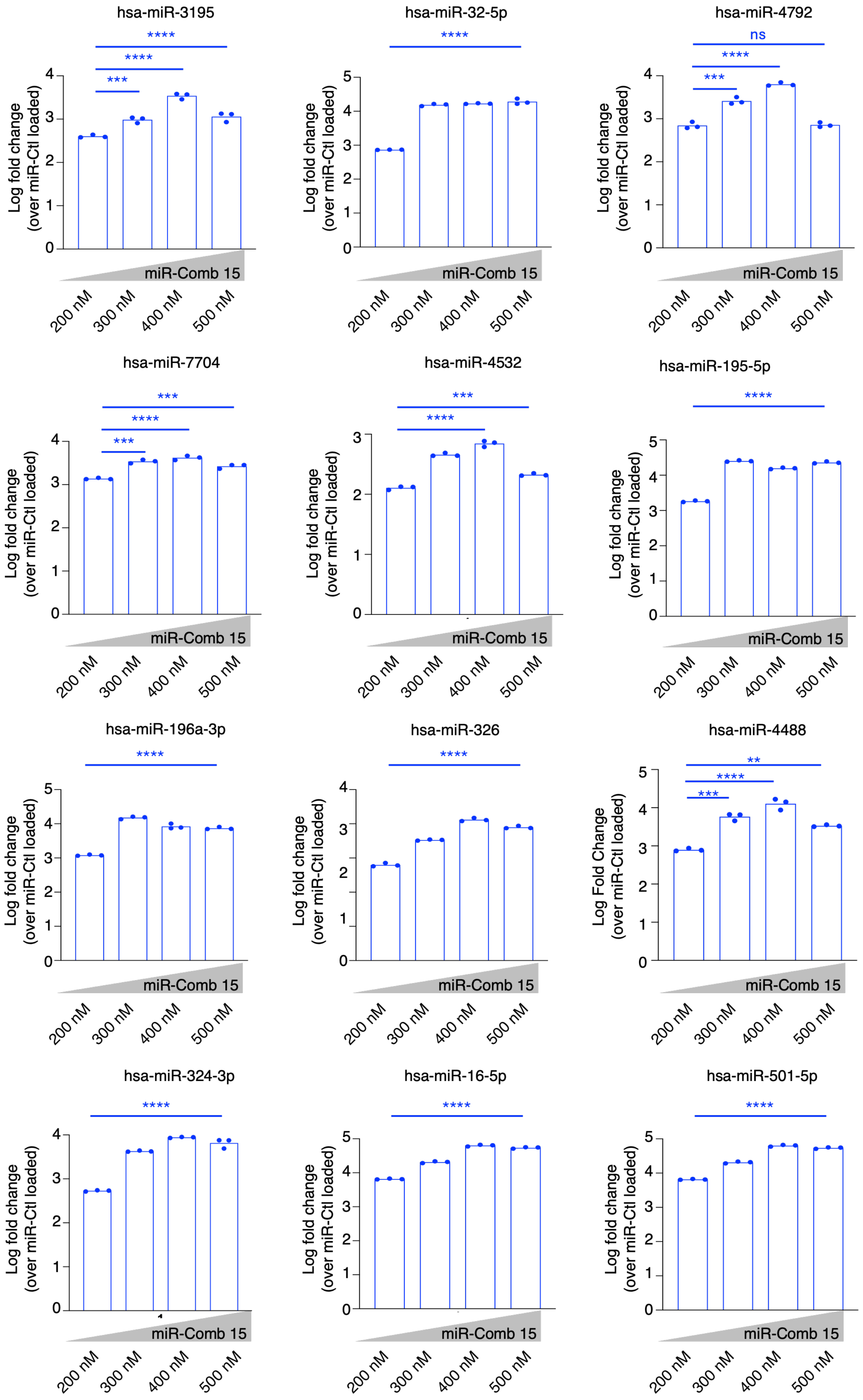
hsa-miR-3195
****
****
***
Log fold change
(over miR-Ctl loaded)
0
1
2
3
4
miR-Comb 15
200 nM
300 nM
400 nM
500 nM
hsa-miR-32-5p
****
Log fold change
(over miR-Ctl loaded)
0
1
2
3
4
5
miR-Comb 15
200 nM
300 nM
400 nM
500 nM
hsa-miR-4792
ns
****
***
Log fold change
(over miR-Ctl loaded)
0
1
2
3
4
miR-Comb 15
200 nM
300 nM
400 nM
500 nM
hsa-miR-7704
***
****
***
Log fold change
(over miR-Ctl loaded)
0
1
2
3
4
miR-Comb 15
200 nM
300 nM
400 nM
500 nM
hsa-miR-4532
***
****
Log fold change
(over miR-Ctl loaded)
0
1
2
3
miR-Comb 15
200 nM
300 nM
400 nM
500 nM
hsa-miR-195-5p
****
Log fold change
(over miR-Ctl loaded)
0
1
2
3
4
5
miR-Comb 15
200 nM
300 nM
400 nM
500 nM
hsa-miR-196a-3p
****
Log fold change
(over miR-Ctl loaded)
0
1
2
3
4
5
miR-Comb 15
200 nM
300 nM
400 nM
500 nM
hsa-miR-326
****
Log fold change
(over miR-Ctl loaded)
0
1
2
3
4
miR-Comb 15
200 nM
300 nM
400 nM
500 nM
hsa-miR-4488
**
****
***
Log Fold Change
(over miR-Ctl loaded)
0
1
2
3
4
5
miR-Comb 15
200 nM
300 nM
400 nM
500 nM
hsa-miR-324-3p
****
Log fold change
(over miR-Ctl loaded)
0
1
2
3
4
miR-Comb 15
200 nM
300 nM
400 nM
500 nM
hsa-miR-16-5p
****
Log fold change
(over miR-Ctl loaded)
0
1
2
3
4
5
miR-Comb 15
200 nM
300 nM
400 nM
500 nM
hsa-miR-501-5p
****
Log fold change
(over miR-Ctl loaded)
0
1
2
3
4
5
miR-Comb 15
200 nM
300 nM
400 nM
500 nM

**Figure S6. RT-qPCR quantification of additional miR-Comb 15 members following dose-dependent loading into HEK 293T-derived EVs.**

RT-qPCR quantification of miRNAs composing the miR-Comb15 signature following pH-gradient loading of HEK 293T-derived EVs with increasing concentrations of miR-Comb 15 (200–500 nM). miRNA abundance is expressed as log fold change relative to miR-Ctl-loaded EVs. A dose-dependent increase in miRNA incorporation was observed, with maximal loading generally reached between 300 and 400 nM, followed by a plateau at higher concentrations. Data are presented as the mean of three independent biological experiments (n = 3). Statistical significance was assessed using an unpaired two-tailed Student's t-test. ns, not significant; *P < 0.01; **P < 0.001; ***P < 0.0001.

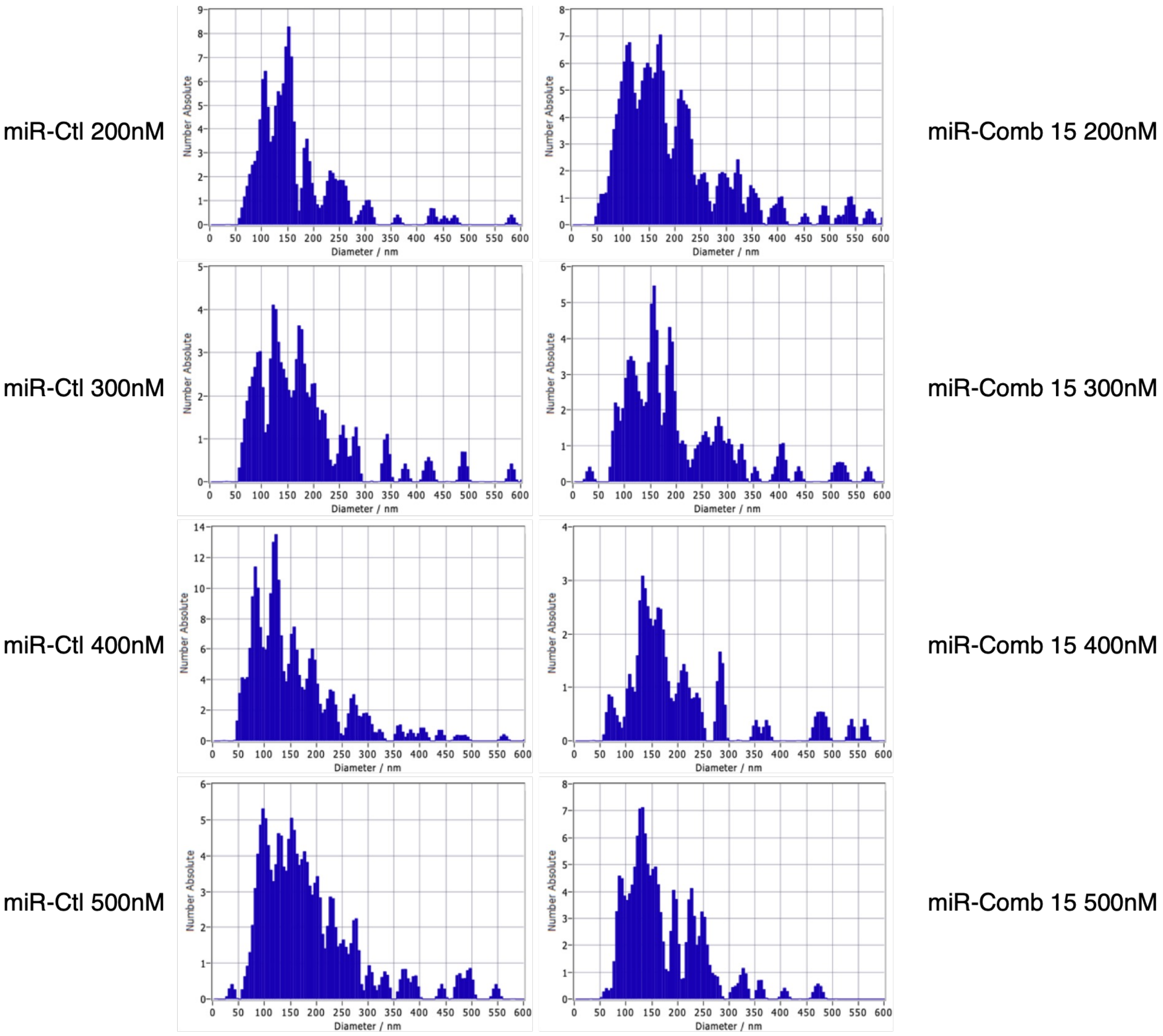


**Figure S7. Particle size distribution of miRNA-loaded HEK 293T-derived EVs.** Nanoparticles tracking analysis (NTA) of HEK 293T-derived EVs following pH-gradient loading with miR-Ctl or miR-Comb 15 at increasing input concentrations (200, 300, 400, and 500 nM). Representative particle size distributions are shown for each loading condition. EVs displayed a predominant size range consistent with small extracellular vesicles, and no major alterations in particle size distribution were observed following miR-Comb 15 loading compared with miR-Ctl-loaded EVs, indicating that the loading procedure did not substantially affect EV size characteristics.

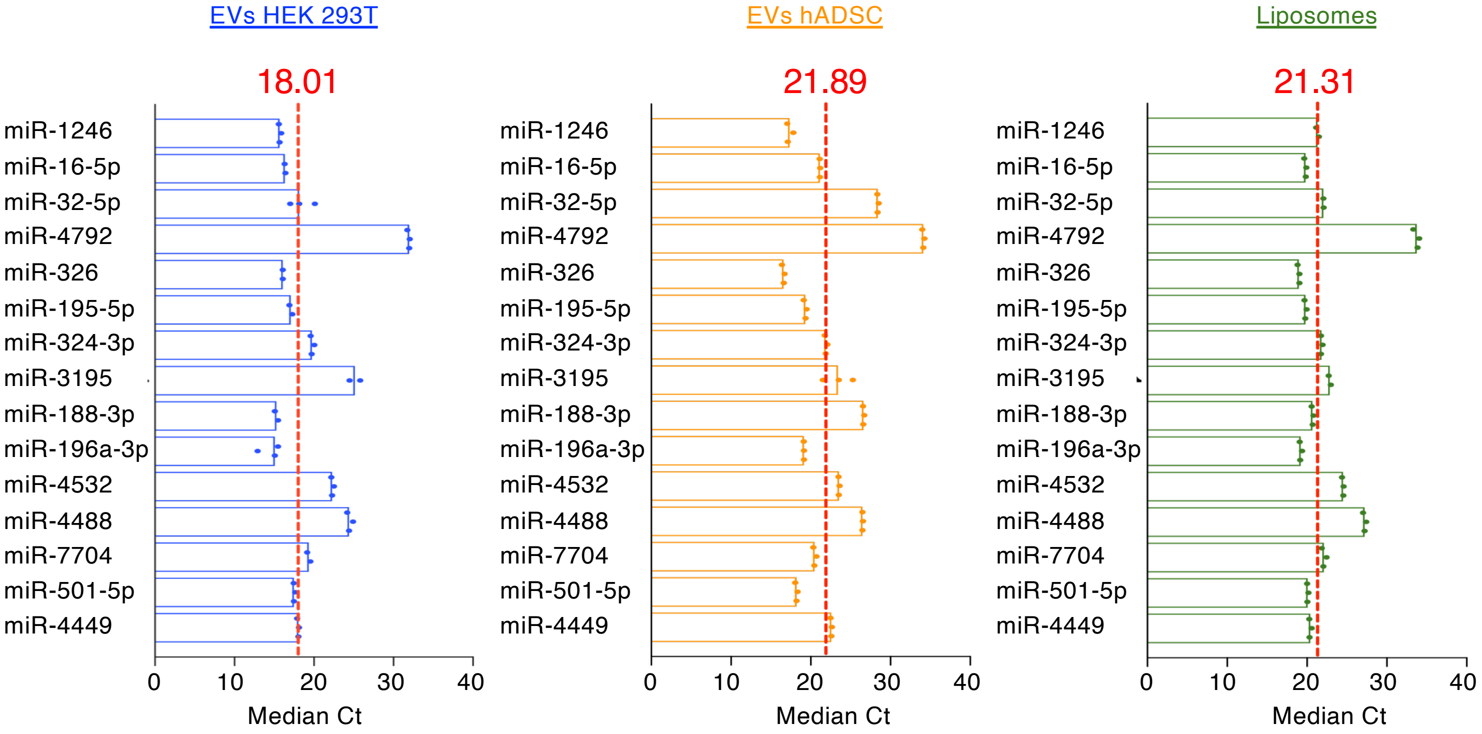


**Figure S8. Median Ct values of miR-Comb15 components following loading into HEK 293T derived EVs, hADSC derived EVs, and liposomes.** RT-qPCR analysis of the 15 miRNAs composing the miR-Comb15 signature following exogenous loading into HEK 293T-derived EVs, hADSC-derived EVs, or synthetic liposomes. Bars represent the median Ct value obtained for each miRNA across independent loading experiments. Red dashed lines indicate the overall median Ct value for each delivery platform. Lower Ct values correspond to higher miRNA abundance. These data provide a comparative overview of miR-Comb15 incorporation across the different delivery systems and complement the loading analyses presented in Figure 6. Data are presented as mean from three independent experiments (n = 3).

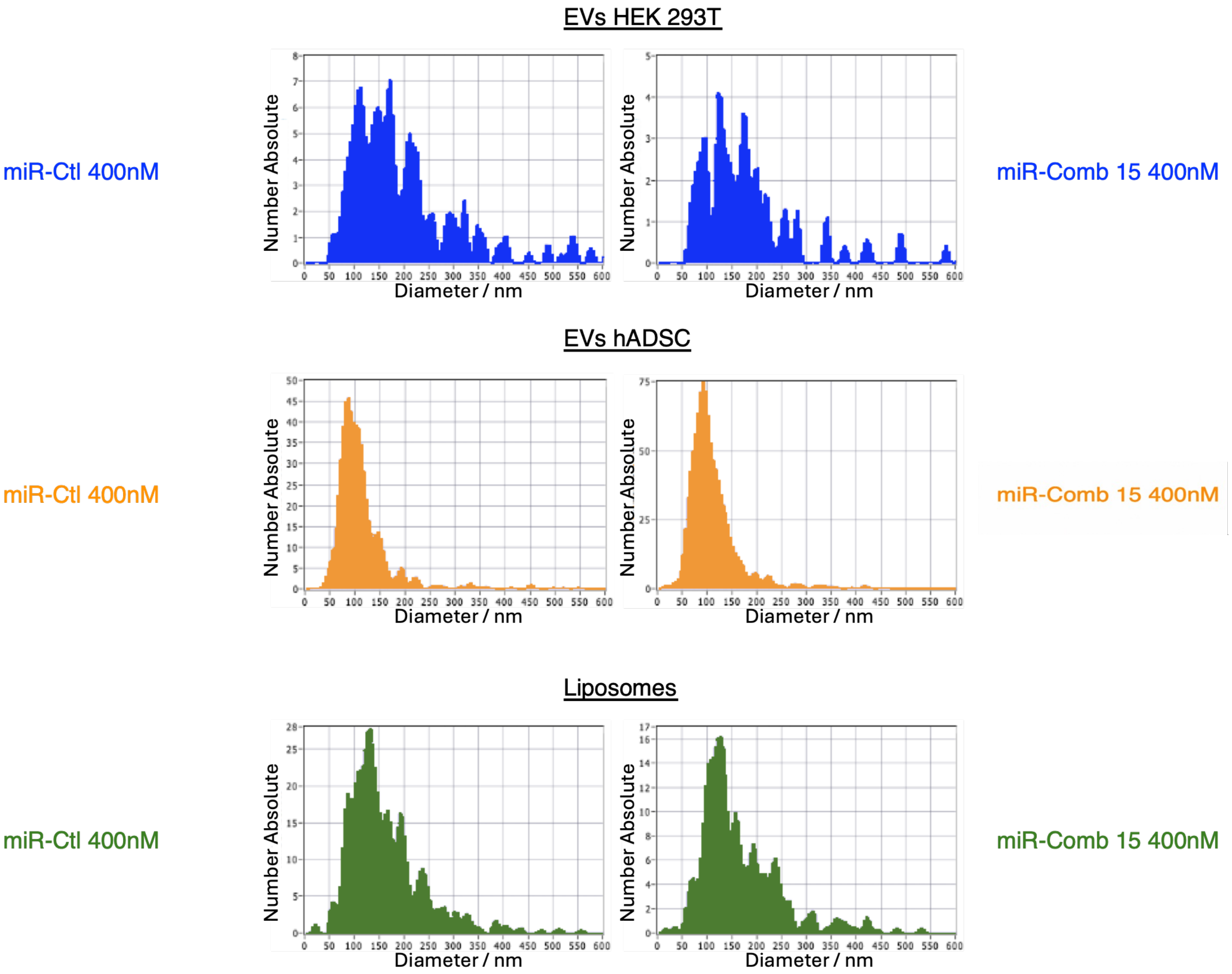


**Figure S9. Particles size distribution of miRNA-loaded delivery platforms.** Representative nanoparticles tracking analysis (NTA) profiles of HEK 293T-derived EVs, hADSC-derived EVs, and liposomes following loading with either miR-Ctl or miR-Comb 15 under the conditions selected for subsequent functional studies. Particles size distributions were determined after loading and purification procedures. Comparable size profiles were observed between miR-Ctl- and miR-Comb15-loaded preparations for each delivery platform, indicating that miRNA loading did not substantially alter particle size characteristics.

| Sequences of miRNAs and corresponding antagomirs used in this study | | | |
|---|---|---|---|
| **Antagomir** | **Sequence (5′→3′)** | **miRNA** | **Sequence (5′→3′)** |
| anti-miRIDIAN Negative Control | UCUGCCGUUUGAUCACGAGCUU | miRIDIAN Negative Control | AAGCCGGUUUGGUAGCUUUAAUU |
| anti-hsa-miR-195-5p | UAGCAGCACAGAAAAUAUUGGC | hsa-miR-195-5p | UAGCAGCACAGAAAAUAUUGGC |
| anti-hsa-miR-1277-3p | UACGUAGAUAUAUAUGUAUUUU | hsa-miR-1277-3p | UACGUAGAUAUAUAUGUAUUUU |
| anti-hsa-miR-501-5p | AAUCCUUUGUCCCUCGGGUGAGA | hsa-miR-501-5p | AAUCCUUUGUCCCUCGGGUGAGA |
| anti-hsa-miR-3960 | GGCGGCGGCGGAGGCGGGGG | hsa-miR-3960 | GGCGGCGGCGGAGGCGGGGG |
| anti-hsa-miR-7641 | UUGAUCUCGGAAGCUAAGC | hsa-miR-7641 | UUGAUCUCGGAAGCUAAGC |
| anti-hsa-miR-33a-5p | GUGCAUUGUAGUUGCAUUGCA | hsa-miR-33a-5p | GUGCAUUGUAGUUGCAUUGCA |
| anti-hsa-miR-629-5p | UGGGUUUACGUUGGGAGAACU | hsa-miR-629-5p | UGGGUUUACGUUGGGAGAACU |
| anti-hsa-miR-324-3p | ACUGCCCCAGGUGCUGCUGG | hsa-miR-324-3p | CCCACUGCCCAGGUGCUGCUGG |
| anti-hsa-miR-188-5p | CAUCCCCUUGCAUGGUGGAGGG | hsa-miR-188-5p | CAUCCCCUUGCAUGGUGGAGGG |
| anti-hsa-miR-4449 | CGUCCCGGGGCUCGCGGAGGCA | hsa-miR-4449 | CGUCCCGGGGCUCGCGGAGGCA |
| anti-hsa-miR-16-5p | UAGCAGCACGUAAAUAUUGGCG | hsa-miR-16-5p | UAGCAGCACGUAAAUAUUGGCG |
| anti-hsa-miR-196a-3p | CGGCAACAAGAAACUGCCUGAG | hsa-miR-196a-3p | CGGCAACAAGAAACUGCCUGAG |
| anti-hsa-miR-32-5p | UAUUGCACAUUACUAAGUUGCA | hsa-miR-32-5p | UAUUGCACAUUACUAAGUUGCA |
| anti-hsa-miR-4488 | AGGGGGCGGGCUCCCGCG | hsa-miR-4488 | AGGGGGCGGGCUCCCGCG |
| anti-hsa-miR-326 | CCUCUGGGCCCUUCUCUCCAG | hsa-miR-326 | CCUCUGGGCCCUUCUCUCCAG |
| anti-hsa-miR-4792 | CGGUGAGCGCUCGCUGUGGC | hsa-miR-4792 | CGGUGAGCGCUCGCUGUGGC |
| anti-hsa-miR-3195 | CGCGCCGGGCCCGGGUU | hsa-miR-3195 | CGCGCCGGGCCCGGGUU |
| anti-hsa-miR-1246 | AAUGGAUUUUUGGAGCAGG | hsa-miR-1246 | AAUGGAUUUUUGGAGCAGG |
| anti-hsa-miR-7704 | CGGGGUCGCGCGCGGACGUG | hsa-miR-7704 | CGGGGUCGCGCGCGGACGUG |
| anti-hsa-miR-188-3p | CUCCCACAUGCAGGGUUUGCA | hsa-miR-188-3p | CUCCCACAUGCAGGGUUUGCA |
| anti-hsa-miR-4532 | CCCCGGGGAGGCCCGGCG | hsa-miR-4532 | CCCCGGGGAGGCCCGGCG |

**Table S1. Sequences of miRNAs and corresponding antagomiRs used in this study.** The table lists the mature miRNA sequences and the corresponding miRIDIAN Hairpin Inhibitors (antagomirs) used for the functional screening and validation experiments. Negative control miRNA and antagomiR sequences are also included.

| Engineered EV preparation | Experiment 1 (%) | Experiment 2 (%) | Mean ± SD (%) |
|---|---|---|---|
| EVs miR-Comb 15 | 2.80 | 3.78 | 3.29 ± 0.69 |
| EVs miR-Ctl | 6.79 | 4.99 | 5.89 ± 1.27 |

**Table S2. Loading efficiency of miRNA incorporation into HEK 293T-derived extracellular vesicles.** Loading efficiency (%) was calculated as the percentage of input miRNA recovered in EV preparations following loading using the 400 nM miRNA solutions employed throughout this study. Values represent two independent experiments quantified by Qubit analysis. SD, standard deviation.
